# ARID1A stabilises non-homologous end joining factors at DNA breaks induced by the G4 ligand pyridostatin

**DOI:** 10.1101/2025.05.10.653238

**Authors:** Susana Simões-Sousa, Noa Amin, Karen A. Lane, Alison Harrod, Malin Pedersen, Mercedes Pardo, Christine Rasetti-Escargueil, Zuzanna Kozik, Katheryn A. G. Begg, Alison Ribeiro, Susana Banerjee, Manuel Stucki, Kevin J. Harrington, Jyoti S. Choudhary, Jessica A. Downs

## Abstract

ARID1A is a subunit of the BAF chromatin remodelling complex that is frequently mutated in cancer. It is challenging to predict how ARID1A loss impacts cancer therapy response because it participates in many different cellular pathways. G quadruplex (G4) binding ligands, such as pyridostatin, have shown anticancer effects, but the pathways and genetic determinants involved in the response to G4 ligands are still not fully understood. Here, we show that ARID1A deficient cells are selectively sensitive to pyridostatin when compared with isogenic controls. Sensitivity to pyridostatin was apparent in ovarian and colorectal cancer cell line models, and *in vivo* studies suggest that G4 ligands hold promise for treating ARID1A deficient cancers. While we find that ARID1A impacts on pyridostatin-induced transcriptional responses, we find that pyridostatin-mediated toxicity in ARID1A-deficient cells is driven by defective DNA repair of topoisomerase-induced breaks. We show that ARID1A-deficient cells are unable to efficiently accumulate non-homologous end joining proteins on chromatin following pyridostatin exposure. These data uncover a role for ARID1A in the cellular response to G4 ligands, and link remodelling to G4 ligand-induced transcriptional and DNA damage responses.

## INTRODUCTION

The SWI/SNF family of chromatin remodellers comprises three complexes: BAF, PBAF, and GBAF, which are defined by their subunit composition.^1^ These complexes share a number of core subunits, and can have either SMARCA4 (BRG1) or SMARCA2 (hBRM) as the catalytic subunit. ARID1A (also called BAF250A) is a subunit that is found only in the BAF complex (Fig. 1A).

**Figure 1.**
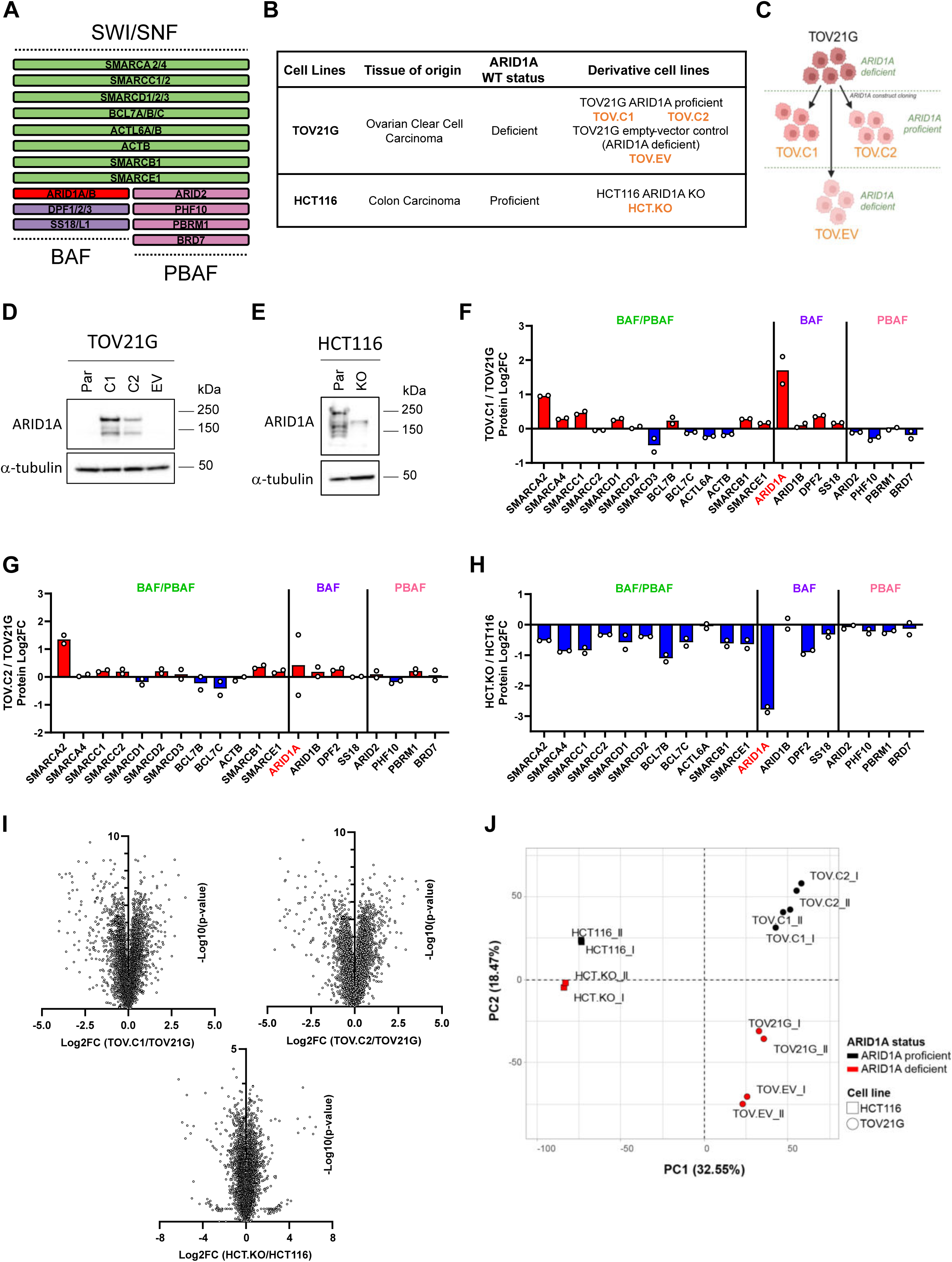
Generation of isogenic ARID1A cell line models. (A) Schematic of the mammalian SWI/SNF chromatin remodelling complexes BAF and PBAF, showing core subunits (green) and complex specific components (purple, BAF; pink, PBAF). ARID1A, a component of BAF complex, is highlighted in red. (B) Summary of isogenic cell lines used in this study. TOV21G, an ARID1A-deficient ovarian clear cell carcinoma cell line, was used to engineer two clones with ARID1A re-expression (TOV.C1 and TOV.C2) and to generate an empty-vector control (TOV.EV). HCT116, an ARID1A-proficient colon carcinoma cell line, was engineered to generate an ARID1A knockout (HCT.KO). (C) Flowchart of TOV21G isogenic cell lines generation. (D and E) Immunoblot analysis of whole-cell extracts showing ARID1A expression in TOV21G cell lines and derivatives (D) and HCT116 parental and knockout cell lines (E). α-tubulin was used as a loading control. (F – H) Quantitative proteomic analysis of the Log2FC (log2 fold change) of SWI/SNF subunits in TOV21G with reconstituted ARID1A cell line TOV.C1 (F) and TOV.C2 (G) compared with parental TOV21G, and in HCT116 ARID1A-knockout (HCT.KO) compared with HCT116 parental cell line (H). Points correspond to individual biological replicates (n = 2). (I) Volcano plots showing differential protein expression in ARID1A-rescued (TOV.C1 and TOV.C2) versus parental TOV21G cell lines and ARID1A knockout HCT116 (HCT.KO) versus parental cell line. (J) Principal component analysis (PCA) of whole proteome data comparing all cell line models used in this study. ARID1A status is indicated by colour (red, deficient; black proficient); different cell lines are distinguished by shape (circle, TOV21G; square, HCT116). See also Figure S1.

ARID1A is involved in multiple pathways that protect genome stability.^1^ It plays a role in both major DNA double strand break (DSB) repair pathways: non-homologous end joining (NHEJ) and homologous recombination (HR). In addition, ARID1A has been implicated in the response to replication stress. ^2,3^ The ARID1A gene is mutated in a wide range of cancer types, including up to 50% of ovarian clear cell carcinomas (OCCC) and around 10% of colorectal cancers ^1^. It is therefore critical to understand when and how each of the functions of ARID1A is used in response to genotoxic agents, particularly those with therapeutic potential.

G quadruplex (G4) structures are secondary structures that can form in guanine-rich single stranded DNA sequences by virtue of the ability of guanines to form G-tetrads that are stabilised by Hoogsteen hydrogen bonding.^4^ These contribute to the regulation of gene expression, and G4s are frequently located in regulatory regions of genes, such as promoters and enhancers.^5^ G4 binding ligands, compounds that can selectively bind to G4 structures, have been shown to impair proliferation or survival of some cancer cells. One way this occurs is through interfering with the expression of oncogenes or other key transcripts that contain G4s in their regulatory regions, thus leading to impaired fitness or viability.^6^

Importantly, another route to impaired cancer cell fitness in response to G4 ligand exposure is through DNA damage. Ligand-stabilised G4 structures lead to DNA double-strand breaks (DSBs) in either a transcription- or replication-dependent manner^7^ and cellular survival following G4 ligand-induced DSBs can be influenced by several cellular pathways.^8–10^ For instance, stabilised G4s can create barriers for the replication machinery, leading to stalled or collapsed replication forks. This outcome can be avoided through the activity of helicases that resolve G4s, such as BLM.^11^ When replication conflicts are not avoided, BRCA1- and BRCA2-dependent homologous recombination (HR) activity has been shown to promote survival following exposure to G4 ligands.^9,12^ Outside of S phase, stabilised G4s have been shown to trap TOP2A on chromatin in a transcription-dependent manner.^10,13^ This results in lesions that rely on TDP2-dependent processing followed by repair via non-homologous end joining (NHEJ).^10^ The clinical exploration of G4 binding ligands is underway, and gaining a deeper understanding of the genetic alterations in cancer cells that makes them vulnerable to G4 ligands will enhance their targeted and effective application.

Here, we generated isogenic cell lines to study ARID1A and found that deficiency leads to increased cellular sensitivity to G4 binding ligands, including pyridostatin (PDS). Sensitivity to G4 ligands was apparent in both ovarian clear cell and colorectal cancer cell line models, and *in vivo* studies suggest that G4 ligands could hold promise for treating ARID1A deficient cancers. We show that the transcriptional response to pyridostatin is influenced by ARID1A, but that the major impact of ARID1A loss on viability following pyridostatin treatment is through DSB repair. Specifically, we find that NHEJ activity is important for the ARID1A-dependent sensitivity to pyridostatin, and we show that ARID1A deficient cells are unable to efficiently assemble non-homologous end joining proteins on chromatin following exposure to pyridostatin. These data show that ARID1A is an important determinant for cellular responses to G4 ligands as a result of its ability to modulate NHEJ repair activities.

## RESULTS

### Generation of isogenic ARID1A cell lines

In order to gain deeper insights into ARID1A biology, we set out to create cell lines with genetic modulation of the ARID1A subunit of BAF (Figure 1A). Because of the prevalence of ARID1A mutations in OCCC, we began by re-expressing ARID1A in the TOV21G cell line, which is an OCCC cell line that has loss of function mutation in ARID1A alleles leading to lack of protein expression. We generated two independently derived clonal cell lines in which we introduced an ARID1A expression construct, referred to hereafter as TOV.C1 and TOV.C2 (Figures 1B and 1C). ARID1A expression was apparent by Western blot analysis (Figure 1D) and immunofluorescence (Figure S1A), with expression levels higher in C1 than C2. As an additional control, we introduced the empty vector (EV) into the TOV21G cell line (TOV.EV; Figures 1B-D and S1A). The ARID1A expressing lines show a modest decrease in proliferation (Figure S1B), but this is also evident in the EV control, suggesting it is not solely due to ARID1A activity and there is no substantial difference in the cell cycle profile of the lines (Figure S1C).

ARID1A is also frequently misregulated in colon cancer, so we made use of a CRISPR-Cas9 knockout (KO) mutation of ARID1A in the HCT116 colon cancer cell line (Figure 1B). We confirmed absence of expression by Western blotting and immunofluorescence (Figure 1E and S1A). Here, we find the absence of ARID1A leads to a modest defect in proliferation (Figure S1B) as well as modest changes to the cell cycle profile (Figure S1C).

To further characterise both cell line models, we performed proteomic analysis using quantitative mass spectrometry. As expected, we found increased ARID1A expression in the TOV.C1 and TOV.C2 cell lines (Figures 1F and 1G). In addition, the levels of several other BAF complex subunits, most notably SMARCA2, were also modestly higher in the ARID1A-expressing lines, likely reflecting increased BAF complex stability (Figures 1F and 1G). In the HCT116 cells, we found that the ARID1A KO cells displayed loss of ARID1A protein expression as well as most other BAF subunits, again consistent with an impact on BAF complex stability when ARID1A is deficient (Figure 1H). Unsurprisingly, ARID1A status impacted on the proteome, with substantial up- and downregulation of a subset of proteins (Figure 1l). Principal components analysis (PCA) of the proteomic data from all cell lines suggests that, while cell line background accounted for the greatest variability between samples, the loss of ARID1A had a substantial impact in both cell lines (Figure 1J).

### ARID1A deficient cells show selective sensitivity to G4 binding ligands

Treatment of cells with the G4 binding ligand pyridostatin (PDS) often leads to impaired fitness.^6^ To test whether this effect is modulated by the presence of ARID1A, we treated our panel of cell lines with PDS and monitored survival. We found that in both cell line models, survival of ARID1A proficient cells was significantly higher than that of ARID1A deficient cells (Figures 2A and 2B). To ensure that the differential vulnerability to PDS was due to ARID1A deficiency, we used siRNA to deplete ARID1A (Figure S2A) and found that reduced ARID1A expression in either reconstituted TOV.C1 cells or HCT116 parental cells leads to PDS hypersensitivity (Figures 2C and 2D). Using an assay to directly monitor cell death, we found that the ARID1A deficient cells have increased levels of cell death following exposure to PDS when compared with the ARID1A proficient cells (Figures 2E and 2F), together suggesting that ARID1A loss leads to increased sensitivity to PDS.

**Figure 2.**
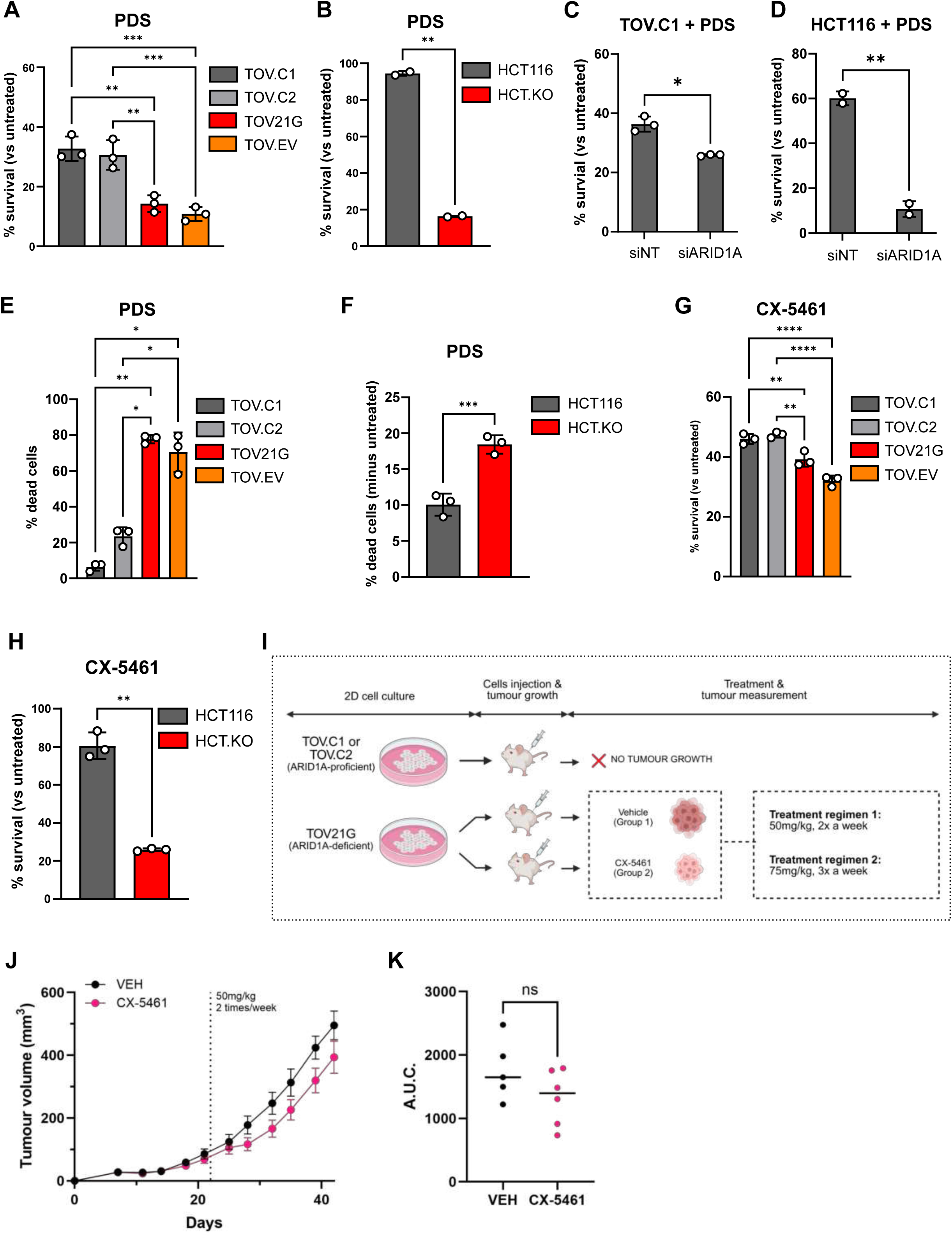
ARID1A deficient cells show selective sensitivity to G4 binding ligands. (A and B) Bar graphs showing percentage survival of isogenic ARID1A cell lines upon treatment with 10 µM PDS for 6 days in (A) TOV21G and (B) HCT116. Viability is expressed as a % of viability in untreated cells. Points correspond to independent biological replicates, n = 3, mean ± SEM, data were analysed by ordinary 1-way ANOVA with Tukey’s test (TOV21G) or n = 2, mean ± SEM, data were analysed by paired t-tests (HCT116), *p<0.05, **p<0.01, ***p<0.001. (C and D) Bar graph showing survival upon treatment with PDS in siRNA-mediated ARID1A depleted cells by SRB assay. Percentage survival of PDS-treated vs untreated is plotted following depletion of ARID1A in TOV.C1 (C) and HCT116 (D) cell lines, upon 6 days-treatment with 10 µM or 20 µM PDS, respectively. Points correspond to independent biological replicates (TOV.C1 n = 3, HCT116 n = 2), mean ± SEM, data were analysed by paired t-tests, *p<0.05, **p<0.01. (E and F) Quantification of dead cells by propidium iodide staining in TOV21G-derived cell lines, following treatment with 10 µM PDS (E) and in HCT.KO versus HCT116 parental, following treatment with 20 µM PDS (F) for 5 days. Points correspond to independent biological replicates, n = 3, mean ± SEM, data were analysed by ordinary 1-way ANOVA with Tukey’s test (TOV21G) or paired t-tests (HCT116), *p<0.05, **p<0.01, ***p<0.001. (G – H) Bar graphs showing percentage survival following 6 days treatment with 8 µM CX-5461 in TOV21G-derived cell lines (G) and following 6 days treatment with 10µM CX-5461 in HCT116 isogenic cell lines. Points correspond to independent biological replicates, n = 3, mean ± SEM, data were analysed by ordinary 1-way ANOVA with Tukey’s test (TOV21G) or paired t-tests (HCT116), *p<0.05, **p<0.01, ***p<0.001, ****p<0.0001. (I) Schematic of the in vivo xenograft experiment. TOV.C1 and TOV.C2 (ARID1A-proficient) or TOV21G (ARID1A-deficient) cells were injected subcutaneously into mice and tumours allowed to form. ARID1A-reconstituted cell lines did not form tumors. Mice bearing tumours derived from TOV21G parental cells were treated with either CX-5461 or vehicle (regimen 1, 50 mg/kg, twice weekly; regimen 2, 75 mg/kg, three times weekly). Figure was generated using BioRender. (J) Tumour growth curve of mice bearing TOV21G xenografts treated with vehicle (VEH) or CX-5461, under treatment regimen 1 (see schematic in (I)). *n*=5 mice for VEH condition, and *n*=6 for CX-5461-treated condition. (K) Area under the curve (A.U.C.) analysis of tumour volume over time from (J). Data were analysed by unpaired t-tests (ns, not significant). See also Figure S2.

ARID1A is found specifically in the BAF complex, but not in the other two highly related SWI/SNF complexes PBAF or GBAF. To see whether this phenotype is shared by other SWI/SNF complexes, we used cell lines with CRISPR-Cas9 mediated KO of PBRM1^14,15^, which is found specifically in the PBAF complex (Figure S2B). These were tested in cell growth assays alongside the isogenic parental RPE1-hTERT cell line. In contrast to our data with ARID1A KO cells, we find no reproducible or significant sensitivity when cells are exposed to PDS (Figure S2C), suggesting that not all SWI/SNF complexes are involved in the cellular response to G4 binding ligands.

CX-5461 is a G4 binding ligand in clinical use.^16^ To see whether the vulnerability of ARID1A deficient cells to G4 ligands has clinical utility, we looked at the response to CX-5461. Like PDS, we find that CX-5461 treatment differentially impacts on ARID1A deficient cells (Figures 2G and 2H). Next, we investigated the response to CX-5461 in mouse models using the ARID1A-deficient ovarian cancer line TOV21G in parallel with the ARID1A-reconstituted cells. We found that ARID1A re-expression resulted in a failure to establish tumours in mice (Figure 2I), consistent with the role of ARID1A as a tumour suppressor.

Mice bearing tumours established from ARID1A-deficient TOV21G cells were treated with vehicle or 50mg/kg CX-5461 twice weekly (Figure 2I). While there was no statistically significant difference in tumour volume or survival between the two groups, there was a trend towards reduced tumour volume in the CX-5461 treated cohort compared with those treated with vehicle alone (Figures 2J and 2K). We therefore tested a higher dose of CX-5461 using 75mg/kg three times weekly. Again, we found a non-significant trend towards reduced tumour volume and increased survival in the CX-5461 treated cohort when compared with vehicle alone (Figures S2D and S2E). These data suggest that there is a potential impact of G4 ligands on ARID1A deficient tumours, but further work is required to determine whether there is clinical value.

### ARID1A loss impacts PDS-dependent transcriptional responses

We next investigated whether the loss of ARID1A influences the transcriptional response to PDS. We performed RNA-seq to analyse the gene expression profile of ARID1A-proficient and ARID1A-deficient TOV cell lines, either without treatment or following 4 h exposure to 10 μM PDS (Figure S3A).

Strikingly, we find that ARID1A has a profound influence on the magnitude of gene expression changes following PDS exposure (Figure 3A). In the presence of ARID1A, there are 1222 genes that are significantly downregulated and 355 that are significantly upregulated in response to PDS. When ARID1A is deficient, these numbers rise to 3954 and 1478, respectively. We find that almost all of the genes whose expression is altered in the ARID1A proficient cells are also significantly changed in the ARID1A-deficient cells (Figure 3B), suggesting that the differences between the samples reflect magnitude of response rather than different pathways.

**Figure 3.**
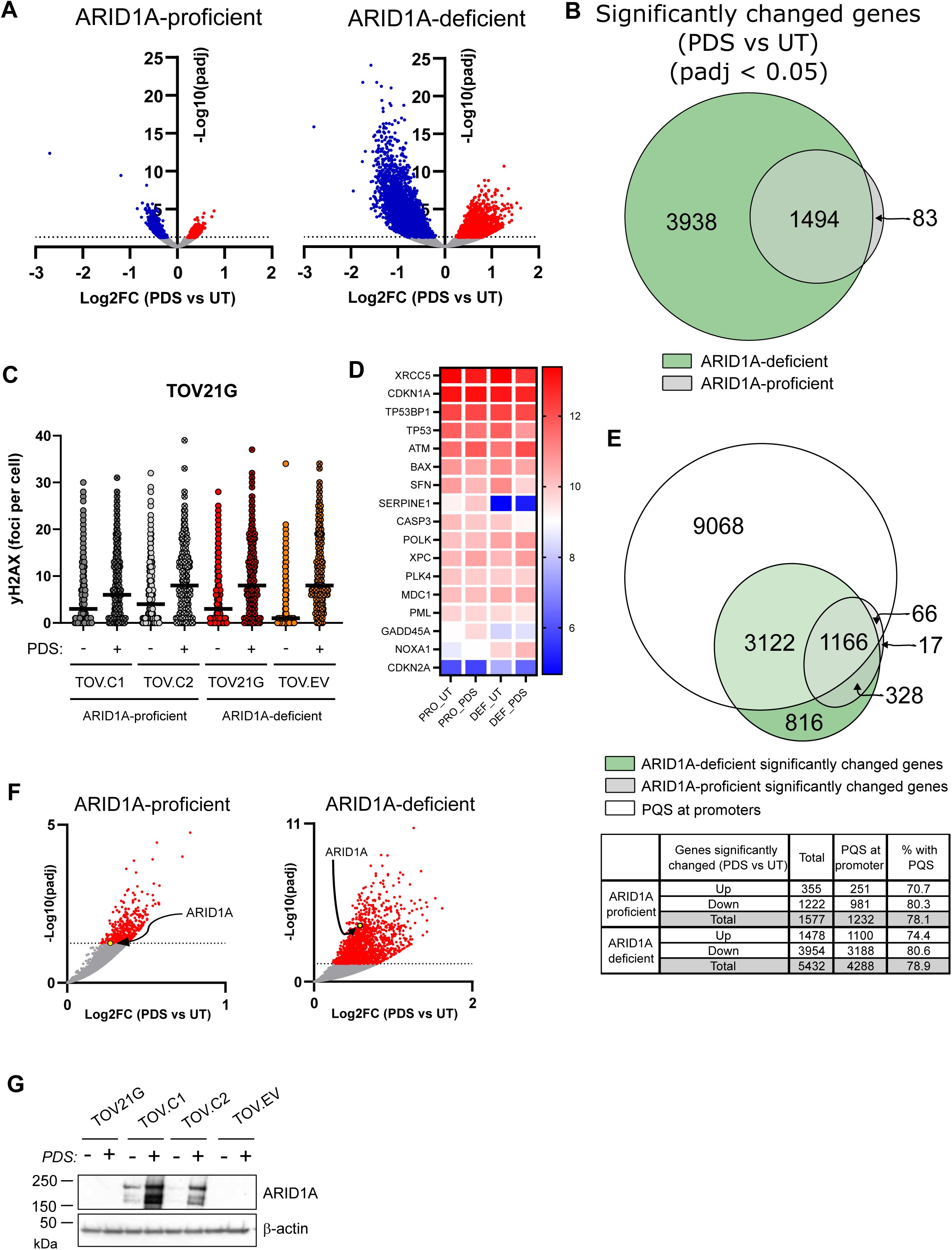
ARID1A loss impacts PDS-dependent transcriptional responses. (A) Volcano plots showing differentially expressed genes in ARID1A-proficient and ARID1A-deficient cells treated with PDS (10 µM, 4 hours) compared to untreated control. Significantly upregulated and downregulated genes are represented in red and blue colours, respectively (adjusted p-value (padj) < 0.05). (B) Venn diagram showing overlap of significantly changed genes (padj < 0.05) upon PDS treatment (10 µM, 4 hours) in ARID1A-deficient and ARID1A-proficient backgrounds. (C) Quantification of γH2AX foci formation in TOV21G isogenic cell lines untreated or treated with PDS (10 µM, 4 hours). (D) Heatmap showing expression of DNA damage induced genes from RNA-seq of ARID1A-proficient and ARID1A-deficient TOV21G isogenic cell lines, untreated or treated with PDS (10 µM, 4 hours). (E) Venn diagram showing the number of significantly altered genes following PDS treatment (10 µM, 4 hours) with potential G quadruplex forming sequences (PQS) at their promoters. Table summarizes the total number of differentially expressed genes, the number of genes with a PQS at their promoter, and the respective percentage for upregulated and downregulated genes in ARID1A-proficient and ARID1A-deficient backgrounds. (F) Volcano plots displaying log2FC of PDS vs UT RNA-seq against the -log10(padj) values in ARID1A-proficient and ARID1A-deficient backgrounds. Significant genes (padj <0.05) are highlighted in red, and ARID1A is highlighted with an arrow. (G) Immunoblot showing ARID1A protein expression response to PDS (10 µM, 6 days) in ARID1A-proficient and ARID1A-deficient cells. β-actin was used as a loading control. See also Figure S3.

PDS exposure leads to DNA double strand breaks (DSBs), and at later time points has been shown to induce inflammatory and p53-dependent transcriptional responses.^17^ Because ARID1A contributes to DSB repair, one potential explanation for these data is that the ARID1A deficient cells have a greater burden of DNA damage, leading to a heightened damage-induced transcriptional response. We therefore tested whether the ARID1A-deficient cells have greater numbers of DNA breaks by looking at γH2AX foci as a readout of DNA DSBs. We used similar conditions as in the RNA-seq analysis (10 μM PDS for 6 h) and find that, as expected, PDS treatment leads to increased γH2AX foci (Figure 3C). Importantly, however, there is no difference between the cell lines (Figure 3C), suggesting that the number of DNA breaks at this time point is relatively similar in the ARID1A-proficient and ARID1A-deficient cell lines. Furthermore, we interrogated the genes with significant PDS-induced changes and find relatively few differential effects on DNA damage responsive genes in any of the datasets (Figures 3D and S3A). Together, these data suggest that differences in DNA damage are not driving the differential transcriptional response to PDS in ARID1A-deficient versus -proficient cells.

Since PDS is not influencing gene expression indirectly via DNA damage, these results suggest it is working through stabilisation of G4 forming sequences that are enriched in gene regulatory elements. If correct, most of the genes whose expression is influenced by PDS will contain G4-forming sequences in their promoters or enhancers. Indeed, we find that almost 80% of the significantly up- or down-regulated genes have predicted G quadruplex forming sequences (PQS) in their promoters (Figures 3E and S3B), which is more than expected by chance since it has been estimated that just over 42% of promoters contain G4 sequences.^18^ This is true in both ARID1A proficient and ARID1A deficient cells (Figure 3E), which suggests that the differential effect of ARID1A is on the magnitude of PDS effects on G4-dependent gene expression of genes with G4s in their promoters. This is likely to be an indirect effect since ARID1A is primarily enriched at enhancers rather than promoters ^19,20^, and the expression of most of these genes is unaffected by ARID1A in untreated cells (Figure S3C).

Interestingly, we noted that ARID1A itself is one of the genes that is upregulated in response to PDS (Figure 3F), and increased protein expression was evident in the ARID1A proficient lines following PDS treatment (Figure 3G). Consistent with our analysis above, ARID1A has a G4 forming sequence in its promoter, and mutation of this sequence was found to reduce ARID1A gene expression.^21^ These data suggest that at early time points, PDS treatment leads to changes of gene expression through modulating G4s in gene regulatory elements and that ARID1A antagonises this response.

### ARID1A-dependent survival following PDS treatment is related to DNA damage responses

The cytotoxicity of PDS is related to DNA damage^10,13^, and ARID1A is known to influence DNA damage responses.^1^ To explore this, we monitored survival over time using live cell imaging. ARID1A deficient TOV21G cells and the two ARID1A-expressing clones were grown in the presence or absence of PDS, and cell fate was monitored over a period of 72 h. We found that a greater number of the ARID1A deficient cells undergo cell death when compared with the ARID1A-expressing lines when treated with PDS (Figures 4A, 4B and S4A). It was evident that, while many ARID1A-expressing cells did not undergo cell death following PDS treatment during this timeframe, they also did not undergo cell division, suggesting that these cells arrested cell cycle progression in response to PDS (Figure 4A). To determine whether the PDS-treated ARID1A expressing cells were viable and able to re-enter the cell cycle, we performed an experiment where PDS was washed out from the media after 4 hours and compared survival with cells treated continuously with PDS (Figure 4C). The survival of the ARID1A-deficient cells was modestly but not significantly different between these treatment conditions (Figures 4B-E). Notably, however, when ARID1A is expressed, survival is significantly greater when cells are transiently treated with PDS compared with continuous treatment (Figure 4B-E).

**Figure 4.**
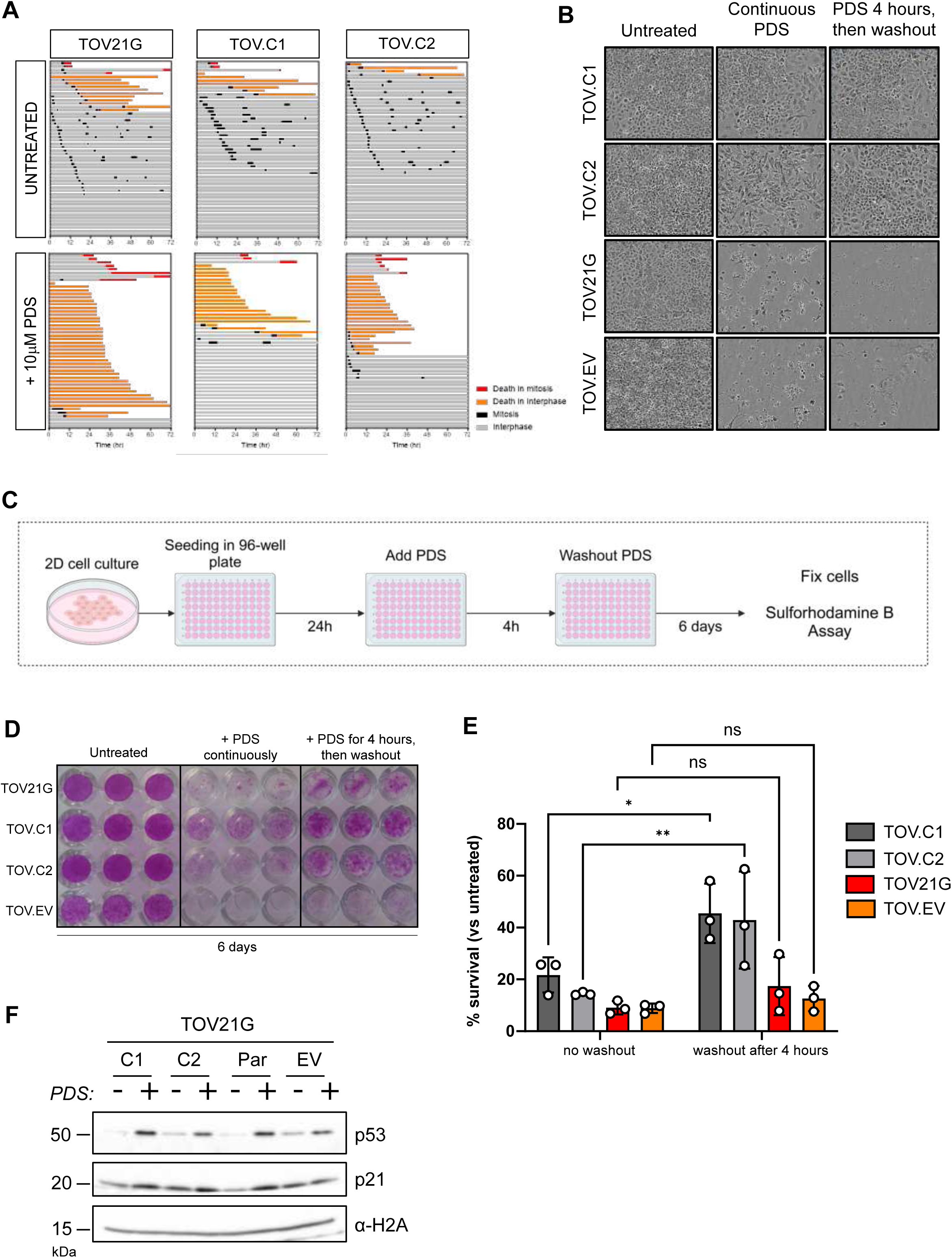
ARID1A-dependent survival following PDS treatment is related to DNA damage responses. (A) Cell fate profiles comparing ARID1A-deficient TOV21G and ARID1A-proficient (TOV.C1 and TOV.C2) following treatment with 10µM PDS for 72 h. Tracks represent individual cells with different colours representing cell cycle events. (B) Representative phase-contrast images of TOV21G-derived cell lines, either untreated or treated with 10 µM PDS for 72 hours or exposed to 10 µM PDS for 4 hours followed by drug washout. Images are from the 72 h timepoint following PDS addition. (C) Schematic of the experimental design used in the viability assays with and without drug washout (in D and E). Figure was generated using BioRender. (D) Representative Sulforhodamine B (SRB)-stained plate showing growth of TOV21G isogenic cell lines across treatment conditions (described in C). Plate was fixed at day 6 after treatment initiation. (E) Quantification of cell survival from SRB assays, normalised to untreated control. Cell survival is shown following 6 days, with cells treated for either 4 hours or continuous treatment with 10 µM PDS. Values show mean ± SEM from three biological replicates, representing three independent experiments. Points correspond to independent biological replicates, n = 3, mean ± SEM, data were analysed by 2-way ANOVA with Šídák’s test, *p<0.05, **p<0.01, ***p<0.001, ****p<0.0001. (F) Immunoblot analysis of p53 and p21 in TOV21G-derived cell lines with or without treatment with 10 µM PDS for 4 hours. α-H2A was used as a loading control. See also Figure S4.

These data suggest that ARID1A proficient cells undergo a transient cell cycle arrest in response to damage caused by PDS. In support of this, we observe a shift in the cell cycle population following PDS treatment (Figure S4B). To look at this more directly, we monitored levels of p53, which is stabilised in response to DNA damage, and its DNA damage upregulated target p21. We found upregulation of both proteins in response to PDS treatment in these cell lines (Figure 4F). This is consistent with previous reports showing a DNA damage checkpoint response to PDS^7,9^, and suggest that in the absence of ARID1A, PDS leads to DNA damage-induced apoptosis, whereas in the presence of ARID1A, cells respond to PDS-induced DNA damage by transiently arresting cell cycle progression.

### ARID1A deficient cells fail to efficiently assemble non-homologous end joining (NHEJ) proteins on chromatin after PDS exposure

The survival and DNA damage response data suggest that ARID1A expressing cells are ultimately able to repair the PDS-induced DNA damage and re-enter the cell cycle, while ARID1A deficient cells are not. Both NHEJ and HR activities have been implicated in mediating survival following PDS exposure^9,10^. To shed light on the repair pathways that might be altered in the absence of ARID1A, we analysed the proteins associated with the chromatin fraction in the TOV21G-derived cell line panel before and after PDS treatment by mass spectroscopy (Figure S5A).

We identified proteins that showed increased abundance in the chromatin fraction in the ARID1A proficient cells upon PDS treatment. Strikingly, many of these proteins were not enriched in the PDS-treated ARID1A deficient chromatin (Figure 5A). When pathway analysis was performed on this group of proteins, we found that the NHEJ pathway was a top hit (Figure 5B). There was very good concordance between the two independently derived ARID1A re-expressing clones (C1 and C2) with NHEJ proteins associating with chromatin post-PDS in both (Figures 5C and S5B). In contrast, these proteins were not substantially enriched in the chromatin fraction of PDS-treated ARID1A deficient TOV21G or EV cells (Figure 5D). These proteins included Ku70 and Ku80, XRCC4, XLF and PAXX (Figure 5E).

**Figure 5.**
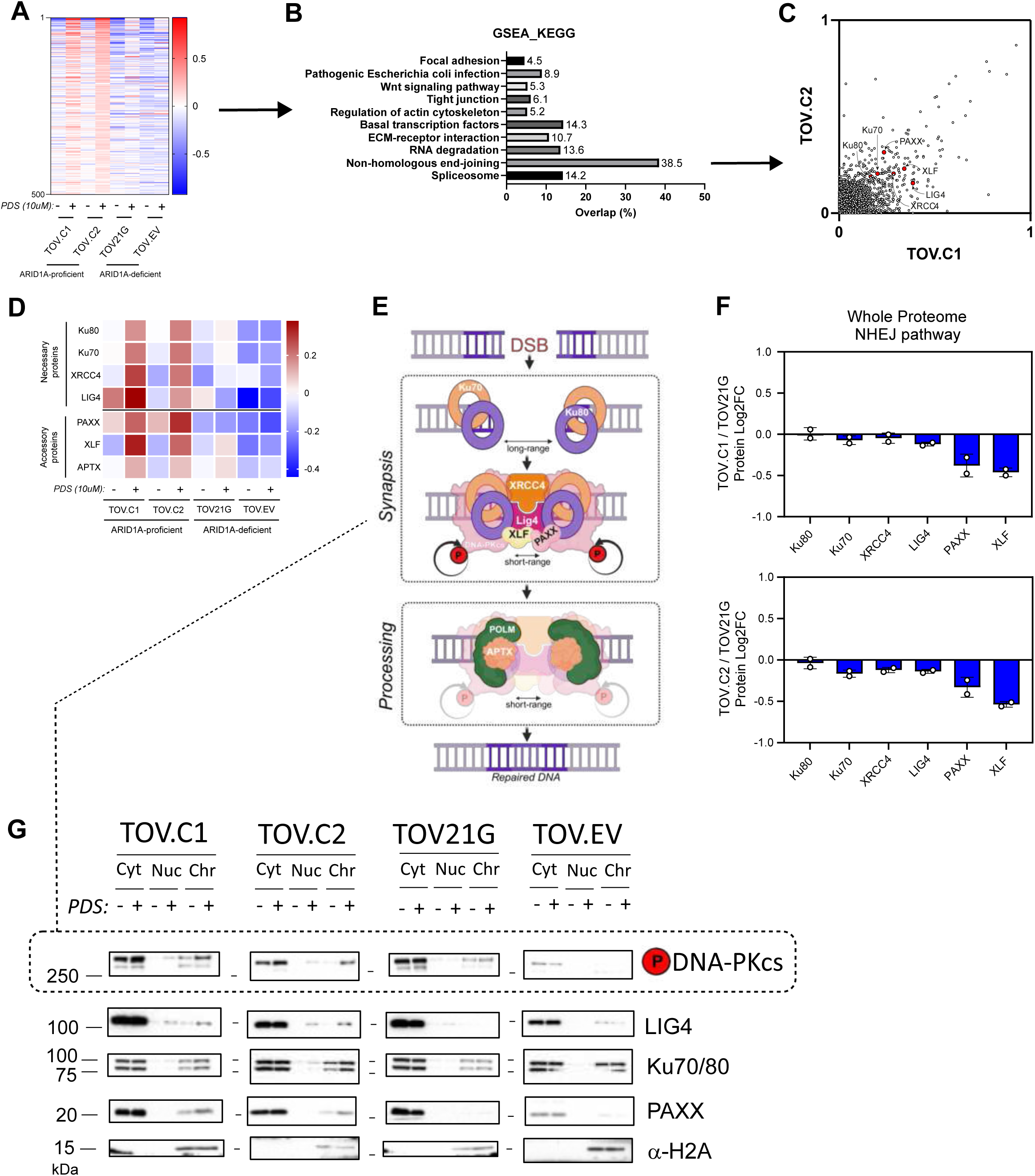
ARID1A deficient cells fail to efficiently assembly non-homologous end joining (NHEJ) proteins on chromatin after PDS exposure. (A) Heatmap of chromatin-associated proteins ordered according to the top 500 most enriched proteins in chromatin in the ARID1A-proficient cell lines (TOV.C1 and TOV.C2) following treatment with 10µM PDS for 4 hours. (B) KEGG pathway enrichment analysis of the 500 proteins represented in (A). Enrichment analysis was done using GSEA and the percentage of proteins in the heatmap (in A) that were in each gene set is plotted. (C) Correlation plot highlighting chromatin-bound levels of a subset of NHEJ proteins in the ARID1A-reconstituted cell lines. (D) Heatmap showing NHEJ factors on chromatin either in untreated conditions or following treatment with PDS (10 µM, 4 hours). (E) Schematic of the NHEJ pathway of double-strand break (DSB) repair. DSB ends are quickly bound by Ku70 and Ku80 proteins, resulting into a long-range synapsis. DNA-PKcs is recruited by Ku proteins, with XRCC4 and LIG4, generating a stable complex (short-range synapsis) that allows the DNA ends to be efficiently re-ligated. Depending on the nature of the broken ends, additional factors (such as POLM, APTX, XLF, and PAXX) contribute to NHEJ repair. Figure was generated using BioRender. (F) Bar graph representing quantitative whole proteome analysis of NHEJ pathway proteins in TOV.C1 and TOV.C2, relative to TOV21G parental cell line. Points correspond to independent biological replicates, n = 2, mean ± SEM. (G) Immunoblots of subcellular fractions (cytoplasmic, nucleoplasmic, and chromatin) prepared from TOV21G-derived cell lines, either untreated or treated with 10 µM PDS for 4 hours. LJ-H2A was used as loading control for the chromatin fraction. See also Figure S5.

We performed a similar analysis in the HCT116 cell line background and find, consistent with the results above, that the ARID1A KO cells have lower levels of core NHEJ proteins associated with chromatin in PDS-treated samples (Figure S5C). Analysis of the proteomes of both TOV21G and HCT116 derived cell lines showed no substantial or consistent difference in the total abundance of NHEJ factors when ARID1A is deficient (Figures 5F and S5D), suggesting that the differential association with chromatin is due to impaired chromatin association, either recruitment or retention, in the absence of ARID1A rather than a deficiency of NHEJ factors.

To investigate this using an orthogonal approach, we performed Western blot analysis of cytoplasmic, nucleoplasmic, and chromatin-bound fractions prepared from the TOV21G cell line panel before and after PDS treatment. We find that there is a detectable increase in chromatin-bound NHEJ proteins after PDS treatment in the two clones with ARID1A re-expression, but this is less apparent in the parental and EV controls (Figure 5G). Using an antibody against phospho-S2056, we similarly find increased phospho-DNA-PK in the chromatin of ARID1A proficient cells (Figure 5G). Similar patterns were observed when we performed the same fractionation analysis in HCT116 parental and ARID1A KO cells (Figure S5E).

### NHEJ activity is important for the repair of DSBs

Together, these data support the notion that there is greater NHEJ activity taking place on chromatin in ARID1A-proficient cells. To test whether NHEJ is functionally important for viability following PDS exposure in these cell lines, we depleted components of the pathway (DNA-PKcs, XLF, and PAXX) and monitored survival. We found that upon depletion of these factors, the ARID1A proficient TOV.C1 cells show reduced viability following PDS survival compared with non-targeting control-depleted cells (Figures 6A, 6B and S6A). In contrast, there was no further reduction in survival when NHEJ factors were depleted in ARID1A-deficient TOV21G cells (Figures 6A, 6B and S6A), suggesting that a major function of ARID1A in promoting survival after PDS is via NHEJ activity.

**Figure 6.**
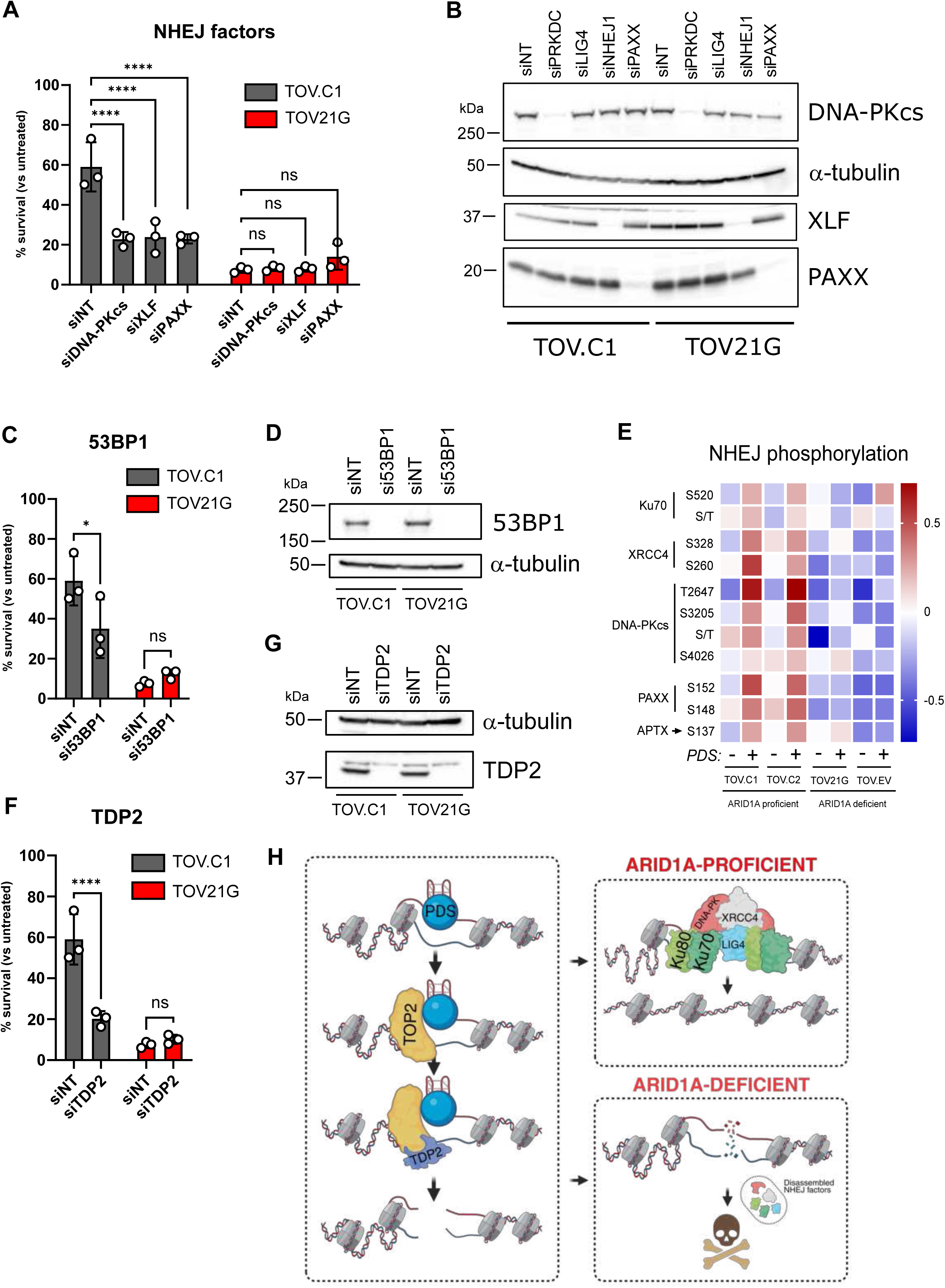
NHEJ activity is important for the repair of PDS-induced DNA damage. (A) Bar graph showing relative survival of PDS-treated ARID1A-proficient (TOV.C1) and ARID1A-deficient (TOV21G) cell lines, following siRNA-mediated knockdown of core NHEJ components (DNA-PKCS, XLF and PAXX). Percentage survival plotted for PDS treated (10 µM, 4 hours) vs untreated cells. Non-targeting siRNA (siNT) was used as a control. Points correspond to independent biological replicates, n = 3, mean ± SEM, data were analysed by ordinary 2-way ANOVA with Dunnett’s test, *p<0.05, **p<0.01, ***p<0.001, ****p<0.0001, ns, not significant. (B) Immunoblots of siRNA depletion of NHEJ components in cells used in (A). α-tubulin was used as loading control. (C) Bar graph showing relative survival of PDS-treated TOV.C1 and TOV21G parental cells following depletion of 53BP1. Percentage survival plotted for PDS treated (10 µM, 4 hours) vs untreated cells. siNT was used as control. Points correspond to independent biological replicates, n = 3, mean ± SEM, data were analysed by ordinary 2-way ANOVA with Fisher’s test, *p<0.05, **p<0.01, ***p<0.001, ****p<0.0001, ns, not significant. (D) Immunoblots of siRNA depletion of 53BP1 in cells used in (C). α-tubulin was used as loading control. (E) Heatmap of phosphopeptides enriched in chromatin-bound NHEJ proteins, in ARID1A-proficient and -deficient cell lines, untreated or upon treatment with 10µM PDS for 4 hours. (F) Bar graph showing relative survival of PDS-treated TOV.C1 and TOV21G parental cells following depletion of TDP2. Percentage survival plotted for PDS treated (10 µM, 4 hours) vs untreated cells. siNT was used as control. Each point corresponds to an independent biological replicate, representing three independent experiments. Points correspond to independent biological replicates, n = 3, mean ± SEM, data were analysed by ordinary 2-way ANOVA with Fisher’s test, *p<0.05, **p<0.01, ***p<0.001, ns, not significant. (G) Immunoblot validation of TDP2 knockdown. α-tubulin was used as loading control. (H) Schematic model. Following TOP2 trapping at PDS-bound G quadruplexes, TDP2 facilitates TOP2 removal, leaving a double-strand break. ARID1A facilitates NHEJ assembly on chromatin, potentially through stabilising short-range synapsis, and enabling effective DSB repair. ARID1A loss impairs NHEJ chromatin assembly or stabilisation, leading to defective DNA repair and ultimately cell death. *All siRNAs were used in parallel in each of the three biological replicates, hence the non-targeting siRNA controls are the same for each set of siRNAs represented in A, C and F*. See also Figure S6.

53BP1 is an upstream regulator of DNA repair pathway choice and promotes NHEJ.^22,23^ We tested the impact of 53BP1 depletion on viability and found very similar patterns as loss of other NHEJ factors (Figures 6C, 6D and S6B), suggesting that 53BP1 is particularly important for PDS responses in ARID1A proficient cells. One possible explanation for these results is that ARID1A interacts with either 53BP1 or NHEJ factors to promote their recruitment or retention on chromatin following PDS treatment. Indeed, ARID1A was recently reported to interact directly with DNA-PKcs.^24^ To test this possibility, we immunoprecipitated ARID1A from extracts prepared from both untreated and PDS-treated cells. We were unable to detect DNA-PKcs, but we did find a weak but reproducible interaction with 53BP1 (Figure S6C). This was not increased by PDS treatment but nevertheless raised the possibility that ARID1A promotes NHEJ by recruiting 53BP1 to chromatin. However, in contrast to NHEJ factors, 53BP1 associated with chromatin did not show a consistent pattern in response to PDS and its chromatin association was not regulated by ARID1A status (Figure S6D). Moreover, we found no difference in 53BP1 foci formation following PDS treatment in ARID1A proficient versus deficient TOV cells (Figure S6E). In HCT116 cells, 53BP1 chromatin association was also unaffected by either PDS or ARID1A (Figure S6F). In addition, the HCT116 ARID1A KO cells have significantly more 53BP1 foci than the ARID1A proficient cells following PDS treatment (Figure S6G). Together, these data indicate that ARID1A is not required for 53BP1 foci formation in response to PDS exposure and suggest that ARID1A is functioning downstream of 53BP1 to facilitate NHEJ activity.

During NHEJ, synapsis follows end binding, thus tethering the two broken ends together to promote repair. Recent work demonstrated that long-range, weakly tethered synapsis is followed by structural remodelling into a more stable, short-range synaptic complex^25^ (Figure 5E). This transition requires DNA-PKcs kinase activity. ^25,26^, and we found that there was a stronger phospho-S2056 of DNA-PK signal in chromatin from ARID1A proficient cells following PDS treatment (Figure 5G). By interrogating the phosphorylation events in the chromatome data, we determined that there was a pronounced enrichment of multiple phosphorylation events of NHEJ factors in the ARID1A proficient, but not deficient, cells following PDS treatment (Figure 6E). These include known DNA damage induced events, such as the well characterised autophosphorylation of DNA-PKcs on T2647.^27–29^ It is possible, therefore, that ARID1A promotes stable short-range synaptic complex formation, thus stabilising NHEJ factors on chromatin at DNA DSBs.

PDS cytotoxicity has been reported to arise through trapping TOP2 intermediates on chromatin. ^10,13^ TOP2 cleavage complexes are removed by TDP2, which allows their repair by NHEJ.^30^ Consequently, if ARID1A promotes survival through NHEJ, loss of TDP2 would prevent ARID1A from doing this. We therefore depleted TDP2 from ARID1A-proficient TOV.C1 cells and find that survival is substantially reduced following PDS treatment, while it has no significant impact on the survival of ARID1A-deficient cells (Figures 6F, 6G and S6H). These data are consistent with a model in which PDS trapped TOP2 must first be removed by TDP2, after which ARID1A promotes NHEJ activity at the DNA breaks (Figure 6H). The absence of ARID1A impairs efficient NHEJ and consequently leads to reduced survival following PDS treatment.

## DISCUSSION

Here, we show that ARID1A is a key determinant of the cellular response to G4 binding ligands such as pyridostatin. We demonstrate that ARID1A influences the transcriptional response of genes with G4-containing promoters to PDS. This includes the ARID1A gene, which is upregulated in response to PDS. However, we find that impaired NHEJ activity is the primary reason for altered viability in ARID1A deficient cells. Interestingly, we can see increased chromatin association by NHEJ factors, but not 53BP1, when ARID1A is present, suggesting that ARID1A functions downstream of 53BP1 to promote NHEJ recruitment or stability on chromatin following DSB formation vias TDP2 processing (Figure 6H). The weak interaction we detect between ARID1A and 53BP1 might indicate that 53BP1 helps to recruit or stabilise ARID1A at sites of PDS-induced damage.

Once localised to PDS-induced DSBs, ARID1A might promote NHEJ indirectly by creating a permissive chromatin environment for NHEJ assembly at DSBs. Given the reported interaction with DNA-PKcs ^24^, It is also possible that ARID1A binds and recruits NHEJ factors to chromatin in the vicinity of PDS-dependent DSBs. Alternatively, or additionally, ARID1A-dependent remodelling activity might promote the transition from long-range to short-range synapsis during NHEJ. Since this transition leads to a more stable association of NHEJ factors with chromatin, a role for ARID1A in promoting this transition would result in an ARID1A-dependent increase in NHEJ factors in the chromatin fraction of PDS-treated cells, in line with what we observe.

Loss of the PBAF specific PBRM1 subunit does not show the same vulnerability to PDS treatment (Figure S2), consistent with the fact that PBAF shows little impact on NHEJ when compared with BAF (for review, see ^1^). It is possible that cell line background plays a role here, but it is likely that the BAF complex, and not PBAF, is important for this activity. In that regard, ARID1B is an orthologue of ARID1A that can be incorporated into the BAF complex instead of ARID1A and has both overlapping and independent cellular roles.^31^ Whether ARID1B-containing BAF complexes contribute to the cellular response to PDS is not yet clear. Unlike findings in other cell line models (for example ^32^), we do not see ARID1B upregulation when ARID1A is deficient in these cell lines (Figure 1), but the ARID1B present in these cells could still promote survival following PDS-induced damage.

One striking finding from this study was the increased magnitude of PDS-induced gene expression changes when ARID1A is absent (Figure 3). There are several potential explanations for this, such as more G4 structures in ARID1A deficient cells or less efficient TOP2 trapping by PDS when ARID1A is present. However, if correct, these possibilities would result in increased DNA DSBs in ARID1A deficient cells, yet we see no substantial differences in γH2AX in this timeframe. Of note, we also found that proteins involved in RNA metabolism, including splicing and degradation, fail to mobilise onto chromatin in the absence of ARID1A following PDS treatment (Figure 5B). Therefore, one potential way by which ARID1A influences gene expression is through degradation, stabilisation, or processing of PDS-regulated transcripts, leading to an apparently muted transcriptional response relative to cells lacking ARID1A. Another, perhaps more appealing possibility is that ARID1A, by virtue of binding to enhancer elements, influences the topological organisation of chromosomes, which might in turn influence the ability of PDS-stabilised G4s to regulate gene expression.

Because G4 ligands are being developed and utilised in the clinic, it is important to understand how common genetic alterations in cancer cells impact on the cellular response to these molecules. Using isogenic cell line models, our findings establish ARID1A deficiency as a vulnerability to the G4 binding ligands PDS and CX-5461. While we demonstrate that impaired NHEJ underlies much of the reduced viability in ARID1A-deficient cells following PDS treatment, it is possible that this is not the sole reason for the differential sensitivity. ARID1A has been implicated in homologous recombination repair of DNA DSBs ^33,34^, and HR deficiency sensitizes cells to these G4 ligands.^9,12^ Moreover, the altered gene expression dynamics could also contribute to the impaired fitness of the ARID1A deficient cells. Given the prevalence of ARID1A mutations in cancers such as ovarian clear cell carcinoma and colorectal cancer, these results provide a good rationale for further evaluation of G4 ligands in these settings.

## RESOURCE AVAILABILITY

### Lead Contact

Further information and requests for resources and reagents should be direct to and will be fulfilled by the lead contact, Jessica Downs.

### Materials availability

All reagents and materials are listed in the key resources table. Cell lines are available at CancerTools.org. All other materials generated in this study are available from the lead contact upon request.

### Data and code availability

Proteomic datasets have been deposited to PRIDE as accession number PXD063556 and transcriptome datasets have been deposited to GEO as accession number GSE295547. These will be publicly available on publication. This paper does not report original code.

## ACKNOWLEDGEMENTS

We thank the Institute of Cancer Research Flow Cytometry facility, Light Microscopy facility, and the Biological Services Unit for support, and we thank the ICR Scientific Computing Team for HPC services. We thank all members of the Downs lab for helpful discussions. The Stucki lab is supported by project grants from the Swiss National Science Foundation (31003A_163141 and 310030_189141) and a grant from the Helmut Horten Foundation. J.S.C. and M.Pa. acknowledge funding from the Wellcome Trust (223745/Z/21/Z), BBSRC (BB/Y004477/1), and ICR Core Funding. This work was further supported by Cancer Research UK (DRCRPG-Jun24/100004; JAD) and the Medical Research Council (MR/W001276/1; JAD).

## AUTHOR CONTRIBUTIONS

S.S., N.A., K.A.L., A.H. and J.A.D. conceived of the study and designed experiments, S.S., N.A., K.A.L., A.H., M.Pa., M.Pe., C.R-E., Z.K., K.A.G.B., and A.R. performed and analyzed experiments, analyzed data, created models, provided resources, S.B., M.S., K.H., J.S.C., and J.A.D. acquired funding and supervised, J.A.D. wrote the manuscript with input from S. S., N.A., K.A.L., and A.H., and all authors contributed to review and editing.

## Key resources table

| REAGENT or RESOURCE | SOURCE | IDENTIFIER |
| --- | --- | --- |
| Antibodies |  |  |
| ARID1A | Merck | HPA005456 |
| γH2AX (Ser139) (20E3) | Cell Signalling | 92845 |
| 53BP1 | Merck | 05-726 |
| Phospho-DNA-PKcs/Phospho-PRKDC (Ser2056) | Cell Signalling | 68716S |
| DNA Ligase IV/LIG4 (D5N5N) | Cell Signalling | 14649S |
| Ku70 (D10A7) | Cell Signalling | 4588S |
| Ku80 (C48E7) | Cell Signalling | 2180S |
| PAXX/C9orf142 | Abcam | ab126353 |
| XLF / NHEJ1 | Abcam | ab189917 |
| TDP2 | Fisher Scientific | 69s87 |
| p53 | Santa Cruz Technology | sc-126 |
| p21 | Santa Cruz Technology | sc-6246 |
| α-H2A | Abcam | ab13923 |
| α-tubulin | Abcam | ab7291 |
| β-actin | Merck | A5060 |
| Goat Anti-rabbit IgG HRP-conjugated | Agilent | P0448 |
| Rabbit Anti-mouse IgG HRP-conjugated | Agilent | P0260 |
| Alexa fluor 488 goat Anti-rabbit | Invitrogen | A11008 |
| Alexa fluor 647 goat Anti-mouse | Invitrogen | A21235 |
| Bacterial and virus strains |  |  |
| XL1-Blue Competent Cells | Agilent | 200130 |
| Biological samples |  |  |
| N/A |  |  |
| Chemicals, peptides, and recombinant proteins |  |  |
| Pyridostatin hydrochloride | Merck | SML2690 |
| CX-5461 | MedChemExpress | HY-13323 |
| Hoechst 33342 trihydrochloride, trihydrate | Merck | H1399 |
| DAPI | Merck | D9542 |
| Critical commercial assays |  |  |
| N/A |  |  |
| Deposited data |  |  |
| PRIDE | https://www.ebi.ac.uk/pride/ | PXD063556 |
| RNA-sequencing | https://www.ncbi.nlm.nih.gov/geo/ | GSE295547 |
| Experimental models: Cell lines |  |  |
| TOV21G | ATCC | CRL-3577 |
| HCT116 | Prof. Manuel Stucki | N/A |
| HEK293TN | Systems Biosciences | LV900A-1 |
| RPE1 | ATCC | CRL-4000 |
| 786-O | ATCC | CRL-1932 |
| Experimental models: Organisms/strains |  |  |
| CD1 | Charles River Laboratories | CrI:CD1(ICR) |
| Oligonucleotides |  |  |
| N/A |  |  |
| Recombinant DNA |  |  |
| pLentipuro-ARID1A | Addgene | #39478 |
| pLentipuro-EV | Addgene | #39481 |
| psPAX2 | Addgene | #12260 |
| MD2G | Addgene | #12259 |
| Software and algorithms |  |  |
| GraphPad Prism | GraphPad | v10.4.2 |
| InkScape | InkScape | v1.4 |
| Incucyte software | Sartorius | v2023A Rev2 |
| SlideBook | 3i | v2024.2 |
| ImageJ | N/A | v1.5.3 or v1.5.4 |
| GNU Image Manipulation Program (GIMP) | N/A | v3.0.2 |
| MetaXpress | Molecular Devices | v6.7.2.290 |
| FlowJo | FlowJo | v10.10.0 |
| R | N/A | v4.3.2 |
| R Studio | N/A | v2024.12.1 |
| ShinyGo | https://bioinformatics.sdstate.edu/go/ | v0.82 |
| BioRender | biorender.com | N/A |

## CONTACT FOR REAGENT AND RESOURCE SHARING

Further information and requests for resources and reagents should be directed to and will be fulfilled by the Lead Contact, Jessica Downs.

## EXPERIMENTAL MODEL AND SUBJECT DETAILS

### Human Cell Lines

Cell lines were authenticated, routinely tested for mycoplasma and cultured at 37°C in a 5% CO_2_ humidified incubator. TOV21G (ATCC), and derived cell lines with reconstituted ARID1A or empty vector, were cultured in base medium consisting of 1:1 mixture of MCDB 105 Medium (Merck) made up in sterile water for cell culture (Merck) supplemented with a final concentration of 1.5g/L sodium bicarbonate (Gibco) and Medium 199 (Merck) supplemented with a final concentration of 2.2g/L sodium bicarbonate, supplemented with 15% FBS (Gibco). HCT116 and their ARID1A KO derivatives (kindly provided by Professor Manuel Stucki, University of Zurich) were cultured in McCoys 5A Modified Media (Gibco) supplemented with 10% FBS, and 1% penicillin/streptomycin (Gibco). HEK293TN (Systems Biosciences) were maintained in Dulbecco Modified Minimal Essential Medium (DMEM)(Merck) supplemented with 10% FBS (Gibco), 200µM glutamax (Gibco), and 1% penicillin/streptomycin (P/S) (Merck). hTERT-RPE1 (RPE1, ATCC) cells were maintained in DMEM/F-12 (Merck) supplemented with 10% FBS, 200µM glutamax (Gibco), 0.26% sodium bicarbonate (Gibco), and 1% penicillin/streptomycin (P/S). 786-O cells (ATCC) were maintained in RPMI 1640 medium (Merck) supplemented with 10% FBS and 1% P/S. PBRM1 knockouts were generated and described in ^15^.

## METHOD DETAILS

### Plasmids

ARID1A re-expression and empty-vector control in TOV21G cell line was performed using the plasmids pLenti-puro-ARID1A (ARID1A) and pLenti-puro-EV (EV) plasmids (gift from the Le-Ming Sheh lab, Addgene plasmids #39478 and #39481, respectively).

### ARID1A re-expression

Plasmids were midiprepped using the Qiagen Plasmid Plus kit as per manufacturer’s protocol and stored at -20°C. They were used to generate viral particles for transfection into ARID1A deficient OCCC cell line, TOV21G. For lentivirus production, 10 µg of ARID1A or EV plasmids were transfected in combination with the lentiviral packaging and envelope plasmids, pMD.2G and psPAX2 using Lipofectamine 3000 according to the manufacturer’s protocol (Thermo Fisher Scientific). Lentiviral production was carried out in HEK293TN. Viral particle-containing medium was harvested at 48 hours and 72 hours post transfection, combined and filtered through 0.45 µm filters, then stored at -80°C. For transduction, viral particle-containing medium was added to parental TOV21G in 10cm^2^ dishes to reach a Multiplicity of Infection (MOI) of 1 with 6 µg/mL polybrene (Merck). Cells were incubated for 24 hours then selected with 1.5 mg/mL G418 (Merck) for seven days. Protein lysates from the resulting cellular pool were screened for ARID1A expression via western blot analysis. Positive clones were single cell sorted into 96-well plates by BD FACSAria™ III sorter (BD Biosciences) then sub-cloned. Positive clones were screened again via western blot analysis and verified via immunofluorescence and mass spectroscopy.

### Immunoblotting

#### Total protein extraction

Total cell protein was extracted by boiling cell pellets in sample buffer (0.35 M Tris pH 6.8, 0.1 g/ml sodium dodecyl sulphate, 93 mg/ml dithiothreitol, 30% glycerol, 50 μg/ml bromophenol blue) to denature and resolved by SDS-PAGE.

#### Chromatin fractionation

For analysis of proteins in different cell compartments, cells were separated into cytoplasm, nucleoplasm and chromatin fractions. For nuclear isolation, cell pellets were incubated on ice with NIB-250 buffer (15mM Tris-HCl [pH=7.5], 60mM KCl, 15mM NaCl, 5mM MgCl_2_, 1mM CaCl_2_, 250mM sucrose) with 1mM DTT, 1x cOmplete™ EDTA-free protease inhibitor cocktail (Roche), 1x PhosSTOP™ phosphatase inhibitor cocktail (Roche), and 0.3% NP-40 for 5 min. Nuclei were isolated by centrifugation at 500 RCF for 5 min at 4°C and supernatant (cytoplasmic fraction) was transferred to a new tube and stored at -20°C. Nuclear pellet was washed with NIB-250, centrifuged at 500 RCF for 5 min at 4°C and resuspended in hypotonic buffer (10mM HEPES [pH=7.9], 3mM EDTA, 0.2mM EGTA) containing 1mM DTT, 1x cOmplete™ EDTA-free protease inhibitor cocktail, and 1x PhosSTOP™ phosphatase inhibitor cocktail, then incubated for 30 min on ice, vortexing every 5-10 mins. Insoluble chromatin fraction was collected by centrifugation for 4 min at 1,700 RCF and supernatant (nucleoplasmic fraction) transferred to a fresh tube and stored at -20°C. The chromatin pellet was resuspended in the appropriate volume of NuPage® 4x LSB (Laemmli Sample Buffer)(Invitrogen) containing 5% β-mercaptoethanol and sonicated using the Bioruptor Pico (Diagenode)(2 x 30 seconds, 30 second gap).

#### Western blotting

Proteins were boiled to denature at 95°C for 5 minutes in 4x LSB containing 5% β-mercaptoethanol, resolved by SDS-PAGE in a Novex™ 4% -20% Tris-Glycine gel (Thermo Fisher Scientific), alongside a Precision Plus Protein Standards (Bio-Rad) for size marker, then electroblotted onto a 0.45 µm nitrocellulose membrane (GE Healthcare Life Sciences).

Membranes were blocked in 5% dried skimmed milk (Marvel) dissolved in TBS buffer containing 0.1% Tween-20 (TBST), and membranes were incubated with primary antibodies dissolved in 5% milk-TBST overnight at 4°C. The following day, membranes were washed three times for 10 minutes with TBST then incubated in HRP-conjugated secondary antibodies diluted in 5% milk-TBST for 1 hour at room temperature. Following three 10-minute washes of the membranes with TBST, proteins were detected using chemiluminescence enhanced Luminata™ Forte Western HRP Substrate (Merck) and imaged on an iBright 750 gel imaging system (Invitrogen). Details of the primary and secondary antibodies used for western blotting in this study are detailed in the Key Resources Table. Western blots were processed for publication using GIMP software (v3.0.2).

### Immunofluorescence

Cells were seeded at appropriate pre-determined densities on 22mm coverslips 48 hours prior to treatment. Cells underwent treatment and were fixed in 4% paraformaldehyde (Electron Microscopy Sciences) in PBS, for 10 minutes at room temperature. After two 4-minute washes with PBS, the paraformaldehyde-fixed cells were permeabilised with 0.5% Triton-X in PBS for 5 minutes at room temperature. Cells were then washed three times for four minutes each with PBS prior to immunostaining. Fixed cells were blocked with 1% BSA-PBS for 1 hour at room temperature, then incubated with primary antibodies diluted in 1% BSA-PBS at the working concentrations, overnight at 4°C. Coverslips were washed three times for four minutes each with PBS, then incubated with secondary antibody and DNA stain 1µg/mL DAPI (Merck) or Hoechst (Merck), indicated in the relevant figures, at the appropriate working concentrations in 1% BSA-PBS, for 1 hour at room temperature and protected from light. After a final three four-minute PBS washes, the slips were mounted onto poly-L-lysine glass slides (Fisher Scientific) with Vectashield mounting medium (Vector Laboratories). The cells were imaged with the Advanced Spinning Disc Confocal Microscope using Slidebook 6 software (3i). All images were obtained with 63x oil lens with z-stacks at relevant intervals. Images were exported as maximum intensity projections for analysis.

### Cell growth analysis

Cells were seeded at 1 x 10^3^ cells/well density in clear bottom black 96-well plates (Greiner Bio-One) and placed in the Incucyte SX5 (Sartorius) equipped with a 10x objective and maintained at 37°C in a humidified 5% CO2 atmosphere, then imaged every four hours, for a total of 72-96 hours. Confluence (% of area covered by cells) was used as a proxy for cell growth. Data was exported and transferred to Graphpad Prism for analysis and presentation.

### Drug treatments

For SRB assays, cell death assays and time-lapse imaging, TOV21G and HCT116 parental and derivative cell lines were treated with 10 µM PDS and 20 µM PDS, respectively, or with 8 µM CX-5461. PDS was dissolved in water, CX-5461 was dissolved in DMSO. Treatment was carried out for 6 days or, for drug washout experiments, for 4 hours then medium containing drug was removed, replaced by normal media and cells were let grow for up to 6 days. PBRM1 KO and matching parental cell lines were treated with a range concentration of PDS (0-50 µM) for 5 days. For cell cycle analysis, microscopy, RNA-seq and proteomics, TOV21G and HCT116 cells lines were treated with 10 µM or 20 µM PDS, respectively, for 4 hours then harvested and analysed. For in vivo experiments, CX-5461 was dissolved in vehicle of 50mM NaH_2_PO_4_.

### Quantitative mass spectrometry and proteomics data analysis

#### Chromatin enrichment

Flash frozen cell pellets were thawed on ice and resuspended in nuclear extraction buffer (15 mM Tris-HCl pH7.5, 60 mM KCl, 15 mM NaCl, 5 mM MgCl2, 1 mM CaCl2, 250 mM sucrose, 0.3 % NP-40, freshly supplemented with 1 mM DTT and Halt Protease and Phosphatase inhibitor (Thermo Scientific) and incubated on ice for 5 min. Nuclei were collected by centrifugation (600 rcf, 5 min at 4°C), washed once with nuclear extraction buffer without NP-40 pelleted again, then re-suspended in pre-chilled hypotonic buffer (3 mM EDTA, 0.2 mM EGTA and freshly supplemented with 1 mM DTT and Halt Protease and Phosphatase inhibitor) and incubated on ice for 30 min to release chromatin. Chromatin was pelleted for 5 min at 1,700 rcf at 4°C cooled centrifuge and subsequently washed twice with hypotonic buffer.

#### MS sample preparation and TMT labelling

Whole cell or chromatin pellets were solubilised using probe sonication in lysis buffer 100 mM triethylammonium bicarbonate (TEAB), 1% sodium deoxycholate (SDC), 10% isopropanol, 50 mM NaCl, 1:1,000 Pierce Universal Nuclease (Thermo Scientific) supplemented with Halt Protease and Phosphatase inhibitor. Protein concentration was measured with the Quick Start™ Bradford protein assay (Bio-Rad) following the manufacturer’s protocol. Protein with an equal contribution from each individual sample were reduced with 5 mM tris-2-carboxyethyl phosphine (TCEP) for 1 hour, followed by alkylation with 10 mM iodoacetamide (IAA) for 30 min, then digested by adding trypsin (Pierce) at final concentration 75 ng/μL to each sample and incubating the samples for 18 hours at RT. For each sample, 100 μg of protein digest was used for phospho-peptides enrichment, while 30 μg was allocated for chromatin and whole-cell proteomics. Phospho-enriched peptides were labelled with TMT-10plex or TMTpro multiplexing reagents (Thermo Scientific) according to the manufacturer’s protocol. Once the samples were combined, SDC was precipitated by adding formic acid (FA) to a final concentration of 2% (v/v), followed by centrifugation at 10,000 rpm for 5 min. The supernatant containing TMT-labelled peptides was dried using a centrifugal vacuum concentrator.

#### High-pH reversed-phase peptide fractionation and liquid chromatography–mass spectrometry analysis

Prior to MS, TMT-labelled peptides were fractionated with high-pH reversed-phase (RP) chromatography using the Waters XBridge C18 column (2.1□×□150□mm, 3.5□μm) on a Dionex UltiMate 3000 high-performance liquid chromatography (HPLC) system. Mobile phase A was 0.1% ammonium hydroxide (v/v), mobile phase B was 100% acetonitrile and 0.1% ammonium hydroxide (v/v). Peptide separation was performed with a gradient elution at 200 μL/min with the following steps: isocratic for 5 min at 5% phase B, gradient for 40□min to 35% phase B, gradient to 80% phase B in 5□min, isocratic for 5 min, and re-equilibrated to 5% phase B. Fractions were collected in a 96-well plate every 42□s to a total of 65 fractions, then concatenated into 12 fractions, dried and reconstituted in 0.1% TFA.

#### Phosphopeptides enrichment

Fractionated peptides were reconstituted in 10 μL of binding solution containing 20% isopropanol and 0.5% formic acid, then loaded onto 10 μL of phospho-peptides enrichment resin (High-Select Fe-NTA Magnetic Phosphopeptide Enrichment Kit, Thermo) pre-washed and conditioned with binding solution in custom-made filter tips fitted onto Eppendorf tubes. Binding was performed at room temperature for 30 min, followed by three washes with 40 μL of binding solution. Washes were performed by centrifugation at 300 g, and the flow-through fractions were collected for total proteome analysis. Phosphopeptides were eluted in three steps using 30 μL of 40% acetonitrile and 400 mM ammonium hydroxide. Both eluted phosphopeptides and flow-through samples were vacuum-dried and stored at −20°C until LC-MS analysis.

#### LC-MS Analysis

LC-MS analysis was performed on the Dionex UltiMate 3000 UHPLC system coupled with the Orbitrap Fusion Lumos or Orbitrap Ascend mass spectrometers (Thermo Scientific). Samples were analysed with the EASY-Spray C18 capillary column (75 μm × 50 cm, 2 μm) at 50 °C. Mobile phase A was 0.1% formic acid and mobile phase B was 80% acetonitrile, 0.1% formic acid. The gradient separation method was as follows: 150 min gradient up to 38% B, for 10 min up to 95% B, for 5 min isocratic at 95% B, re-equilibration to 5% B in 10 min, for 10 min isocratic at 5% B. Precursors between 375 and 1500□m/z□were selected with a mass resolution of 120□000, automatic gain control (AGC) of 4 × 105, and IT (injection time) of 50 ms, with the top speed mode in 3 s for high collision dissociation (HCD) fragmentation with a quadrupole isolation width of 0.7 Th (Thomson unit). The collision energy was set at 35% (TMTpro) or 38% (TMT-10plex), with AGC at 1 × 105□and IT at 86 ms. The samples were analysed on an Orbitrap Fusion Lumos had following settings: A Dionex Ultimate 3000 system and mass spectrometer (Thermo Scientific) were used for data acquisition. From each fraction 10□μl was injected onto a C18 trapping column (Acclaim PepMap 100, 100 μm × 2 cm, 5 μm, 100 Å) at a 10 μL/min flow rate. The samples were subjected to a low-pH gradient elution on a nanocapillary reversed phase column (Acclaim PepMap C18, 75□μm × 50□cm, 2□μm, 100□Å) at 45°C. Mobile phases A and B were 0.1% formic acid and 80% acetonitrile, 0.1% formic acid respectively. The separation was performed at 300 nl/min flow rate and 90□min gradient from 5% to 38% phase B followed by 10□min up to 95% phase B, isocratic for 5□min at 95% B, re-equilibrated to 5% phase B in 5□min, and isocratic for 10□min at 5% phase B. MS1 scans were collected with mass resolution of 120,000, automatic gain control of 4□×□105, and injection time of 50□ms. Precursor ions (Top Speed mode, 3 s) were fragmented with collision-induced dissociation (CID) with a quadrupole isolation width of 0.7 Th (Thomson unit). Collision energy was set at 35%, with AGC at 1□×□104 and IT at 50□ms. Quantification was obtained at the MS3 level with higher-energy collisional dissociation (HCD) fragmentation of the top 7 most abundant CID fragments isolated with synchronous precursor selection (SPS). Quadrupole isolation width was set at 0.7 Th, collision energy was applied at 65%, and the AGC setting was at 1□×□105 with IT at 105□ms. The HCD MS3 spectra were acquired for the range 100-500 m/z with a resolution of 50,000. Targeted precursors were dynamically excluded for further fragmentation for 45□s with 7 ppm mass tolerance. Phosphopeptide samples were analysed with an HCD method at the MS2 level with CE 38%, AGC 1 × 105 and max IT 105 ms. The samples analysed on Orbitrap Ascend had following settings: MS1 scans were performed over a mass range of m/z 400–1600 at 120,000 resolution in the Orbitrap, with standard AGC settings and automatic injection times. Ions with charge states +2 to +6 were included. Dynamic exclusion was set to 45 seconds with a repeat count of 1, a ±10 ppm mass tolerance, and isotopes were excluded from further analysis. MS2 spectra were acquired in the ion trap using a Turbo scan rate, with 32% HCD collision energy and a maximum injection time of 35 ms. Real-time database searching against Homo sapiens (canonical and isoforms) was conducted using the Comet search engine, considering tryptic peptides with a maximum of 1 missed cleavage. Static modifications were set for carbamidomethylation of C (+57.0215 Da) and TMTpro labelling on K and N-termini (+304.207 Da). Variable modifications included deamidation of N/Q (+0.984 Da) and oxidation of M (+15.9949 Da), allowing up to 2 variable modifications per peptide. A close-out feature was enabled, limiting to 4 peptides per protein. SPS10-MS3 scans were performed on selected precursors using the Orbitrap at 45,000 resolution with 55% HCD collision energy, a 200% normalized AGC target, and a 200 ms maximum injection time. Data were collected in centroid mode with single micro-scan acquisition.

#### Database search and protein quantification

The SequestHT and Comet search engines were used to analyse the acquired spectra in Proteome Discoverer 2.4 and 3.0 (Thermo Scientific) for protein identification and quantification. For chromatin and whole cell proteomes, the precursor mass was set to 20 ppm and fragment mass tolerance was 0.5 Da. Spectra were searched for fully tryptic peptides with maximum 2 missed-cleavages. TMTpro at N-term/Lys and Carbamidomethyl at Cys were defined as static modifications. Dynamic modifications included oxidation of Met and Deamidation of Asn/Gln. For phospho-enriched peptides, the precursor mass was set to 10 ppm and fragment mass tolerance was 0.02 Da. TMT-10plex or TMTpro at N-term and Carbamidomethyl at Cys were defined as static modifications, while dynamic modifications included oxidation of Met, Deamidation of Asn/Gln, TMTpro or Acetyl at Lys residue. Peptide confidence was estimated with the Percolator node. Peptide FDR was set at 1% and validation was based on q-value and target-decoy database search. Spectra were searched against reviewed UniProt human protein entries. The reporter ion quantifier node included a TMT quantification method with an integration window tolerance of 15 ppm and integration method based on the most confident centroid peak at the MS3 or MS2 level. Only unique peptides were used for quantification, considering protein groups for peptide uniqueness. Peptides with average reporter signal-to-noise >3 were used for quantification. The data was normalised to total loading at the proteome level, while for the respective phospho-proteome the data was corrected for loading for phosphorylated peptides only. Relative abundances were calculated by dividing normalised protein/peptide abundances by the average abundance of all TMT channels per biological replicate. PCA plot was generated in R by combining relative normalised abundances HCT116 and TOV21G datasets into a single dataset using dyplr (v1.1.4) and tidyverse (v2.0.0) and correlations were calculated and plotted using FactoMineR (v2.11).

### Cell cycle analysis

Cells were trypsinised, washed with PBS and fixed in 70% ice-cold ethanol 96 hours after treatment with or without 10 µM PDS (Merck). Samples were stored overnight at -20°C. Fixed cell pellets were washed once in PBS then incubated in PBS containing 5 µg/mL propidium iodide (Merck) and 100 µg/mL RNAse A (Merck) for 30 minutes at 37°C and protected from light. Analysis of the stained cells were carried out on the BD LSR II flow cytometer or the BD FACSymphony™ A5 flow cytometer with 20,000 single cell events recorded per sample after gating. Data analysis was performed using FlowJo v10.10.0 software.

### Sulforhodamine B (SRB) assays

Cells were seeded in 96-well plates at the appropriate densities to reach 70 - 80% confluence after 5-6 days in the absence of treatment. This was at 15,000 cells/mL (1,500 cells/well) for TOV21G cell lines, 10,000 cells/ml (1,000 cells/well) for the HCT116, RPE1, HEK293, and 786-O cell lines. Cells were allowed to adhere overnight before addition of drug. Cells were allowed to grow for 5-6 days, after which they were fixed by adding 100 µl 10% TCA (Merck) and incubated at 4°C for 24 hours. Plates were then washed five times with water and allowed to completely dry before staining with 100 µl 0.057% SRB (Merck) in 1% acetic acid (Fisher Scientific) and incubated for 1 hour room temperature. Unbound dye was removed by washing with 1% acetic acid, and the plate was allowed to dry completely. Protein-bound dye was solubilised in 100 µl of 10mM Tris-HCl (pH 9.5) for 10 minutes at room temperature. The optical density measurements at 564 nm were quantitated using the SpectraMax Microplate Absorbance Reader (PerkinElmer). Cell growth was expressed as a percentage of the absorbance measured in the treated wells compared with the control wells.

### Cell death assay (Celigo)

Cells were seeded in 96-well plates at the appropriate densities to reach 70 - 80% confluence after 6 days in the absence of treatment. Cells were allowed to adhere overnight before treatment with PDS and incubated for 5 days. 30 minutes prior to analysis on the Celigo (Nexelcom Bioscience), 20 µl media containing propidium iodide (Merck) and Hoechst to a final concentration of 2 µg/mL each was added to well. The proportion of dead cells was defined as the number of PI-positive cells as a percentage of total (Hoechst-positive) cells.

### RNA-seq

For RNA-sequencing, pellets were harvested by scraping cells in ice-cold PBS following by centrifugation at 7,500 x g for 10 minutes. Three independent biological replicates were used for TOV21G parental cells (n=3), and two independent biological replicates of each ARID1A complemented cell line (TOV.C1 and TOV.C2), which were combined for n=4 in total for analysis purposes. Pellets were resuspended in 500µL TRIzol reagent (Ambion, Thermo Fisher Scientific) and total RNA was extracted using a Direct-zol RNA miniprep kit (Zymo Research) according to the manufacturer’s protocol. RNA concentration and quality was confirmed with the High Sensitivity RNA ScreenTape (Agilent), using the Agilent 4150 Tapestation. Library preparation and sequencing was performed by Novogene Corporation Ltd. Novogene NGS Stranded RNA Library Prep Set was used to generate 250-300bp insert strand specific libraries, and ribosomal RNA was removed using TruSeq Stranded Total RNA Library Prep. 50 million 150bp paired-end reads were sequenced on an Illumina NovaSeq 6000. Fastq reads were checked using FastQC (v0.11.9) (Andrews, S. (2010). FastQC: A Quality Control Tool for High Throughput Sequence Data [Online]. http://www.bioinformatics.babraham.ac.uk/projects/fastqc/) and trimmed using TrimGalore (v0.6.6) (Krueger F, Trimgalore (2023), GitHub repository, https://github.com/FelixKrueger/TrimGalore). Residual ribosomal RNA reads were removed using Ribodetector with -e norRNA setting (v0.2.7)^35^ and strandedness was detected using RSeQC (v4.0.0)^36^. Reads were aligned to the T2T-CHM13v2 genome using STAR alignment software (v2.7.6a)^37^ and reads mapping to genes were quantified using HTSeqCount ^38^. Differential analysis of gene expression was calculated in R using DESeq2 (v1.38.3)^39^. Area-proportional venn diagrams with ellipses were generated using eulerr package v7.0.2 (eulrr Larsson J (2024). eulerr: Area-Proportional Euler and Venn Diagrams with Ellipses). Heatmaps displaying scaled regularised log transformed read counts for each RNA-seq replicate were plotted using ComplexHeatmap v2.20.0^40^, with columns split by condition and rows split by k-means clustering. Gene IDs for each cluster were extracted using dplyr v1.1.4 (Wickham H, François R, Henry L, Müller K, Vaughan D (2023). _dplyr: A Grammar of Data Manipulation) for pathway analysis. Pathway enrichment was calculated using Hallmark signature gene sets ^41^ on ShinyGo ^42^. Potential quadruplex-forming sequences (PQS) in the T2T-CHM13v2 genome were identified using pqsfinder v2.24.0 ^43^ using default settings. Promoter coordinates (hsEPDnew version 006 ^44^) were lifted over from hg38 to T2T coordinates using the UCSC liftOver command line tool. Genes with PQS at promoters were defined using the bedtools (v2.29.2) window tool as genes with promoters within 100bp of a PQS.

### Tumour growth and in vivo treatments

All procedures involving animals were approved by the Animal Welfare and Ethical Review Board at the Institute of Cancer Research in accordance with National Home Office Regulations under the Animals (Scientific Procedure) Act 1986.

For experiments involving tumour growth, TOV21G cells were expanded, then trypsinised and filtered through 40µm cell strainers. 5×10^6^ cells were mixed with Corning Matrigel^TM^ matrix basement membrane (Fisher Scientific) at a 2:1 ratio (cells:Matrigel) in a total volume of 100µL, and were injected subcutaneously in the right flank of 9-12-week-old female CD1 nude mice (Charles River Laboratories). Tumours were allowed to grow for 3-4 weeks and thereafter mice were randomised based on tumour volume (∼100mm^3^) before start of treatment. The number of mice per treatment group is indicated in the figure legends. Mice were treated with CX-5461 or vehicle solution by oral gavage at the indicated times and doses detailed in figure legends. Tumours were measured using callipers twice weekly and tumour volumes calculated using the formula length x width x height (mm) x 0.5236 and data plotted in GraphPad Prism. Data from tumour growth are plotted as mean±SEM and area under curve (A.U.C) was calculated from start of inoculation. Mouse body weights were measured twice weekly.

### High-throughput fluorescence microscopy

TOV21G and HCT116 parental and derivative cell lines were seeded in 96-well plates at the density of 10,000 cells per well. 24 hours later, cells were treated with 10 µM and 20 µM PDS, respectively, for 4 hours, then fixed with 4% paraformaldehyde (Electron Microscopy Sciences) for 10 minutes at room temperature. After three rounds of 4-minute washes with 1xPBS, cells were permeabilised with 0.2% Triton-PBS, for 10 minutes at room temperature. After additional three rounds of 4-minute washes with 1xPBS, cells were blocked with 1% BSA at room temperature for 30 minutes then incubated overnight at 4°C, in the relevant primary antibodies, either ⍰H2AX or 53BP1, both diluted 1:500 in 1% BSA. Cells then underwent three 4-minute washes with 1xPBS and were subsequently incubated with the relevant secondary antibodies (anti-rabbit green 488 for ⍰H2AX and anti-mouse red for 53BP1). After an hour incubation in the secondary antibodies at room temperature, cells were washed three times for 4 minutes in 1xPBS. 200uL PBS was added to each well and cells were analysed using an ImageXpress Microconfocal Spinning Disk system (Molecular Devices) to detect ⍰H2AX and 53BP1 foci, and foci number were quantified using MetaXpress software.

### Cell fate profiling

To measure proliferation, apoptosis induction, and to perform cell fate profiling, 1×10^4^ cells were seeded per well in μclear 96 well plates (Greiner Bio-One) and IncuCyte Kinetic Caspase-3/7 Apoptosis Assay Reagent (Sartorius) added at 1:2000 concentration. Cells were then imaged using an IncuCyte SX5 (Sartorius) equipped with a 10x objective and maintained at 37°C in a humidified 5% CO2 atmosphere. Phase contrast and fluorescence images (3 images per well) were collected every 10 min, using confluence and fluorescence as proxies for proliferation and apoptosis, respectively. Apoptosis was quantitated by measuring green fluorescence object count in 3 images per well. Image sequences were then exported in MPEG-4 format and individual cells were analysed manually to generate cell fate profiles. Timing data were imported into GraphPad Prism for statistical analysis and presentation. Note that 0 hr on the fate profiles represents when imaging started.

### RNA interference

For RNAi-mediated inhibition, a cell suspension of 250,000 cell/ml suspension was prepared in media without antibiotics and mixed with transfection reagent consisting in 1:1 mixture of 120nM siRNA diluted in Opti-MEM® media (Life-Technologies) and 100nM Lipofectamine™ RNAiMAX Transfection Reagent (Thermo Fisher Scientific) diluted in Opti-MEM® media. Each cells/transfection reagent mixture was transferred to a 6cm dish and incubated at 37°C in a 5% CO_2_ humidified incubator for 24h. Knock-down was confirmed by immunoblotting. ON-TARGETplus siRNA pools targeting ARID1A, DNA-PKcs, XLF, PAXX, 53BP1 and TDP2 were obtained from Dharmacon. Non-targeting control was a mixture of Mission siRNA Universal Negative control #2 (Sigma) and siRNA Negative control (Eurogentec).

### Co-immunoprecipitation (CO-IP)

Cell pellets of TOV21G-proficient cell line (TOV.C1) or HCT116 cells (untreated and treated with 10 or 20µM pyridostatin, respectively, for 4 hours) were lysed in 50□mM Tris-HCl pH 8, 300 mM NaCl, 0.2% Igepal, 1 mM EDTA, containing 1 mM DTT and Halt protease and phosphatase inhibitors (Pierce). Lysates were then diluted to 50□mM Tris-HCl pH 8, 150 mM NaCl, 0.1% Igepal, 1 mM EDTA. Anti-53BP1 (Milipore 05-726) and normal mouse IgG (Millipore) (2□µg) were coupled to 25 µl of Protein G Dynabeads (Thermo) according to manufacturer’s instructions. Antibody-coupled beads were incubated with cell lysates containing 0.5-1 mg of protein for 1–2□h at 4□°C. Beads were then washed three times with 10 mM Tris-ClH pH 8, 150 mM NaCl, 0.1% Igepal. Immunoprecipitates were eluted in 1x LDS (Life Technologies) at 70 °C for 10 min. Proteins were separated in 3-8% NuPAGE Tris-Acetate gels (Life Technologies) and transferred to nitrocellulose membranes. Immunodetection was performed according to standard procedures using anti-53BP1 (Abcam ab21083, 1:2000) and anti-ARID1A (Sigma HPA005456, 1:1000) antibodies, followed by incubation with HRP-labelled Protein A (GE Healthcare). Membranes were developed with ECL Prime (Cytiva) or Immobilon ECL Ultra Western HRP Substrate (Millipore) and chemiluminescence was captured with an iBright imager (Invitrogen) or on X-ray film (Hyperfilm ECL, Cytiva).

### Software & Statistical analyses

The number of biological repeats for each experiment and statistical analyses used are indicated in the Fig. legend. Graphs were generated and statistical analyses performed using GraphPad Prism (v10.4.2). Microscopy images were analysed using ImageJ (v1.5.3 or 1.5.4), CellProfiler (v4.0.7) or MetaXpress software (v6.7.2.290), and visualized using GraphPad Prism. Omics data were visualised using the indicated packages in R (v4.3.2) and R Studio (v2024.12.1). Figures were generated using Inkscape Software. Some graphics were generated using BioRender (biorender.com), indicated in figure legends.

---

**Supplemental Figure 1.**
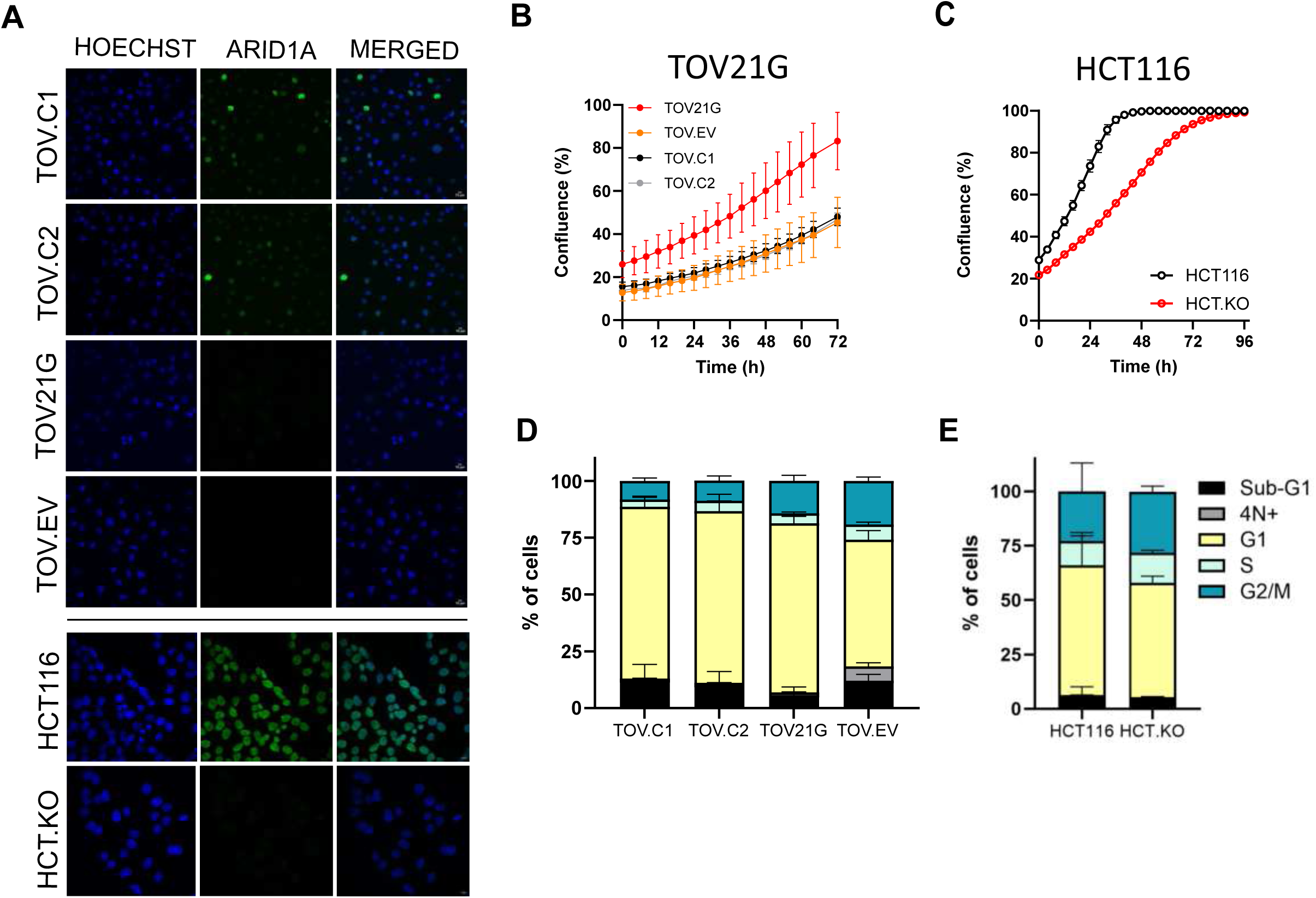
ARID1A expression and its impact on cell proliferation and cell cycle profiles. (A) Immunofluorescence staining of ARID1A (green) and Hoechst (blue) in TOV21G and HCT116 cell lines and derivatives. (B) Real-time confluence measurement of TOV21G (left) and HCT116 (right) isogenic cell lines, over 72 and 96 hours in untreated conditions, respectively. (C) Cell cycle distribution assessed by flow cytometry using DNA content profiling of TOV21G (left) and HCT116 (right) isogenic cell lines in untreated conditions. Data represents independent biological replicates, n = 3, mean ± SEM.

**Supplemental Figure 2.**
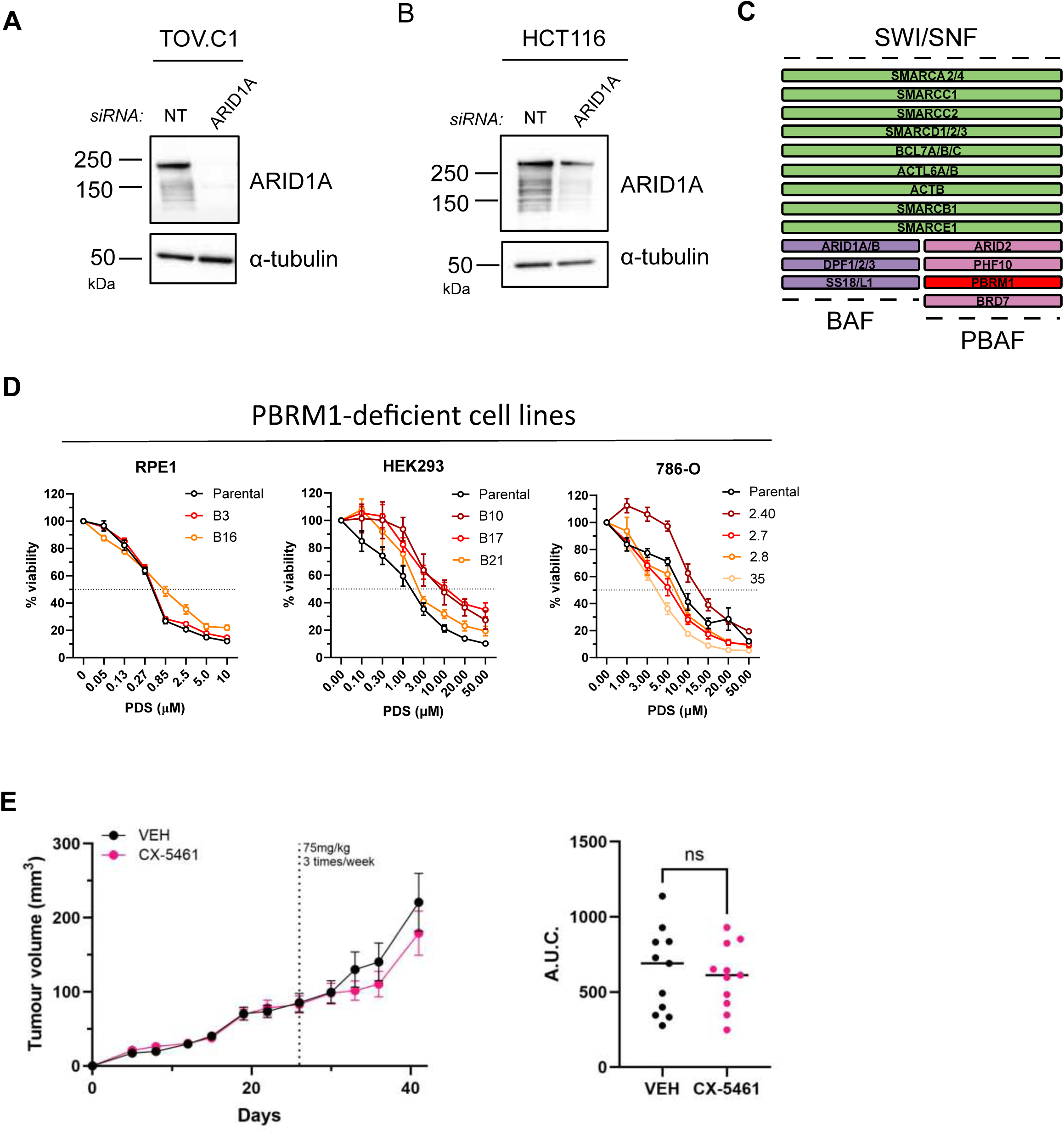
Impact of PBRM1 modulation in the response to G4 stabilisation and *in vivo* response of ARID1A-deficient tumours to treatment with clinical G4 ligand. (A) Immunoblot showing ARID1A siRNA-mediated knockdown efficiency in ARID1A-proficient cell lines. α-tubulin was used as loading control. (B) Schematic of the mammalian SWI/SNF chromatin remodelling complexes BAF and PBAF, highlighting PBRM1, an exclusive component of PBAF, in red. (C) Dose-response curves showing sensitivity to PDS in PBRM1-deficient cell lines derived from RPE1, HEK293 and 786-O cell lines. Data represents independent biological replicates, n = 3, mean ± SEM. (D) Tumour growth curve of mice bearing TOV21G xenografts treated with vehicle (VEH) or CX-5461 under Treatment Regimen 2 (see schematic in Figure 2I). *n*=11 mice for each condition. (E) Area under the curve (A.U.C.) analysis of tumour volume over time from (D). Data were analysed by unpaired t-tests (ns, not significant).

**Supplemental Figure 3.**
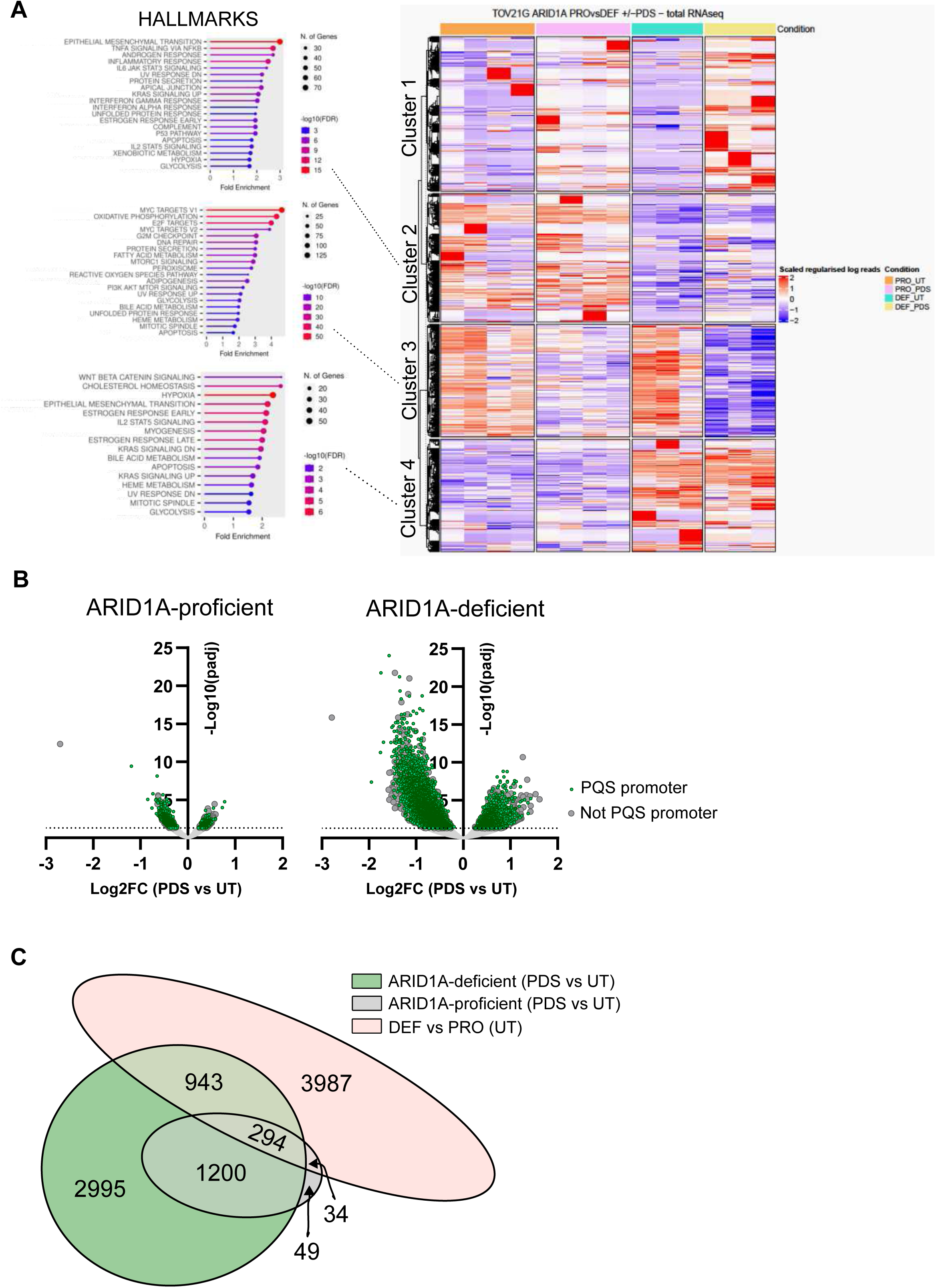
ARID1A status differentially impacts transcriptional responses to G4 ligands. (A) Heatmap showing the scaled regularised log-transformed read counts of the whole transcriptome of ARID1A-proficient and -deficient TOV21G isogenic cell lines, untreated or treated with PDS (10 µM, 4 hours), with samples split by condition and genes split into 4 clusters using k-means clustering. Hallmarks gene set enrichment analysis was carried out on these clusters, using ShinyGO v0.82. Cluster 1 did not have any significantly enriched gene sets. (B) Volcano plots showing differential gene expression induced by PDS treatment (10 µM, 4 hours), in ARID1A-proficient and -deficient cells (as in Fig 3A), highlighting significantly changed genes with promoter proximal to PQS (green) or promoters without PQS (grey). (C) Venn diagram comparing significantly changed genes upon PDS treatment (10 µM, 4 hours) in ARID1A-deficient (green) and -proficient (grey) TOV21G isogenic cells, compared with significantly changed genes in untreated ARID1A-deficient versus ARID1A-proficient TOV21G cell lines (pink).

**Supplemental Figure 4.**
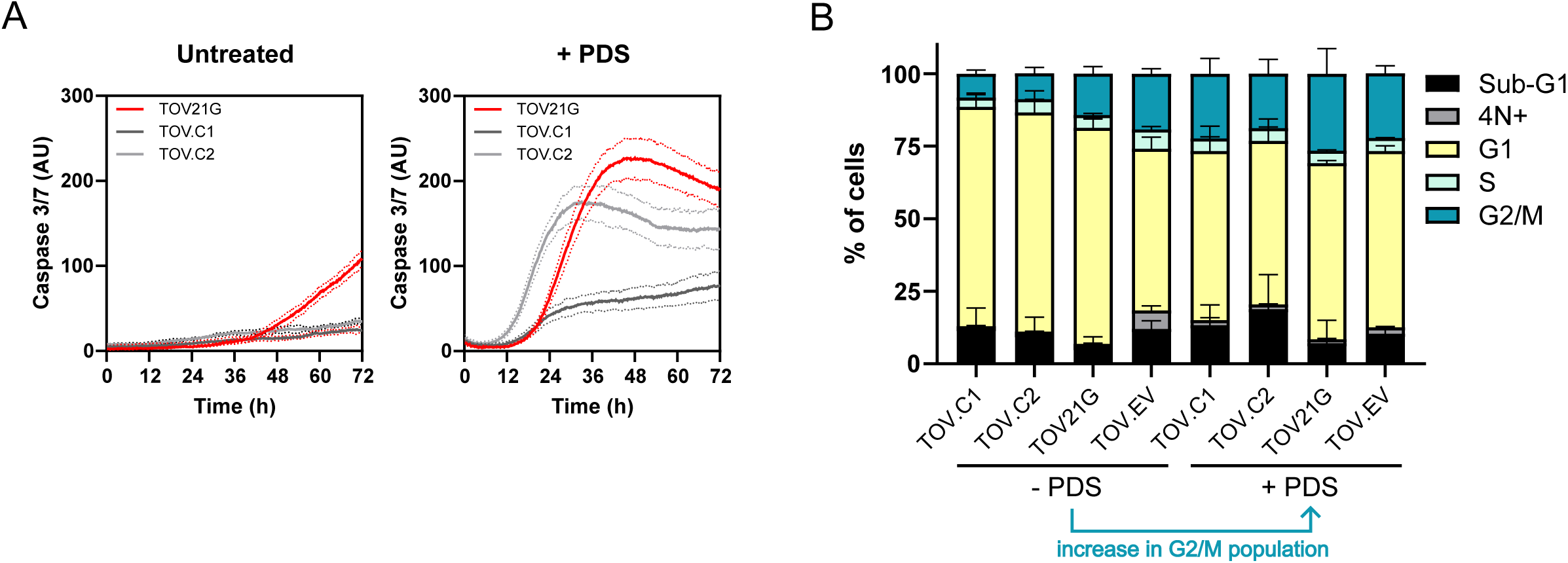
ARID1A loss promotes caspase 3/7 activation, proliferation defects and cell cycle disruption following PDS exposure. (A) Real-time measurement of caspase 3/7 activity in TOV21G, TOV.C1 and TOV.C2 cells over 72 hours in untreated conditions (left) or following PDS treatment (10 µM; right). Data represents technical replicates (n = 4, AU=arbitrary units). (B) Cell cycle distribution in TOV21G-derived cell lines under basal and PDS-treated conditions (10 µM, 4 hours), assessed by flow cytometry measuring DNA content. Data represents independent biological replicates (n = 3). *Note that untreated samples are the same represented in Fig.S1C, which were run in parallel with the PDS treated samples represented in this figure*.

**Supplemental Figure 5.**
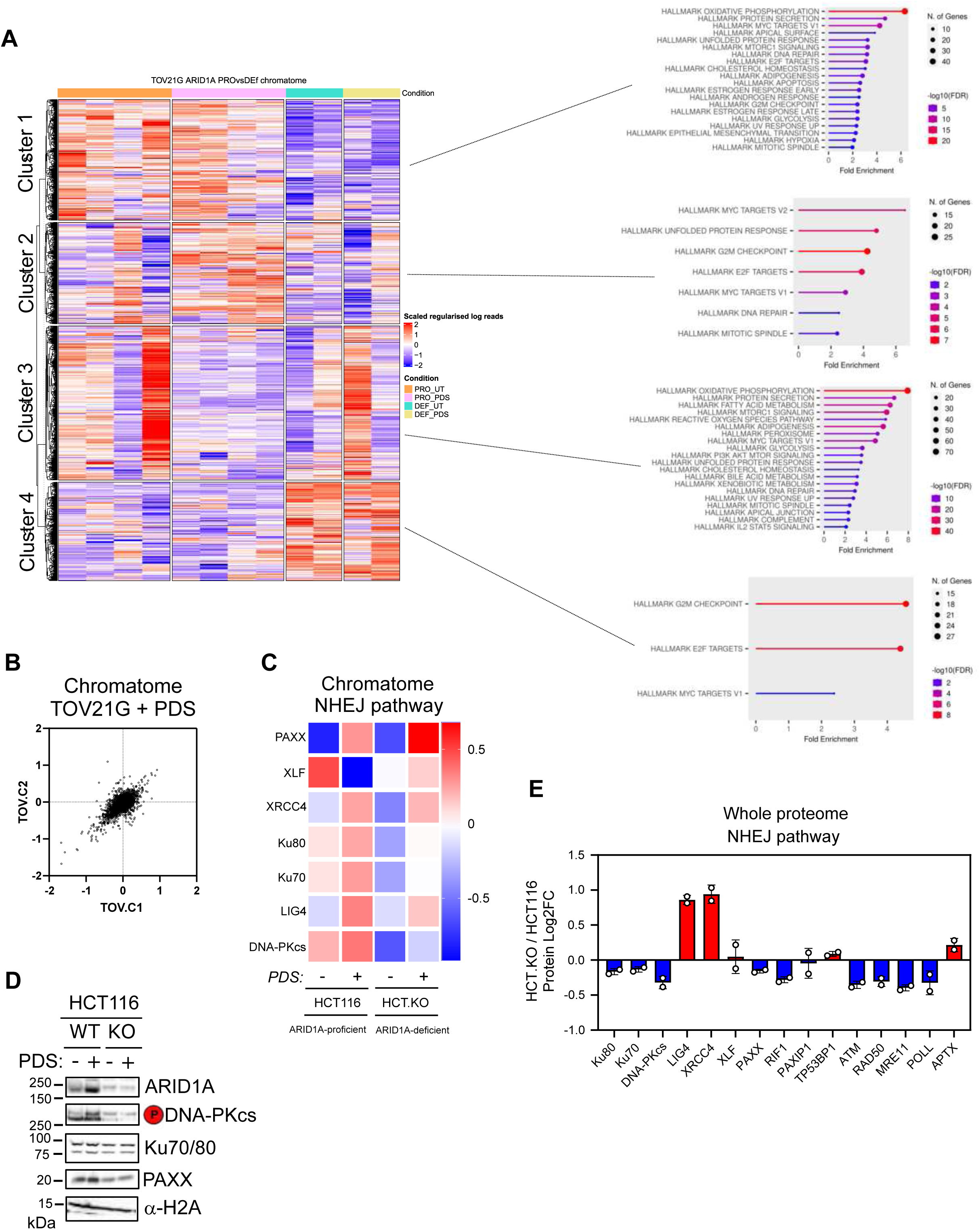
ARID1A status impacts protein chromatin association patterns following PDS exposure. (A) Heatmap of the log2-centred abundance of all chromatin-bound proteins across ARID1A-proficient and -deficient TOV21G cell lines, untreated or treated with PDS (10µM, 4 hours). Hierarchical clustering reveals distinct chromatin recruitment patterns in response to ARID1A status and PDS exposure. (B) Scatterplot comparing chromatin-associated proteome between TOV.C1 and TOV.C2 following PDS treatment (10 µM, 4 hours). (C) Heatmap of NHEJ components from chromatin-associated proteome in HCT.KO versus HCT116 parental cells. (D) Whole proteome quantification of NHEJ components in HCT.KO versus HCT116 parental cells, untreated or treated with PDS (20 µM, 4 hours). Points correspond to independent biological replicates, n = 2, mean ± SEM. (E) Immunoblot analysis of chromatin fractions from HCT116 parental and KO cells, untreated or treated with PDS (20 µM, 4 hours). α-H2A was used as a loading control.

**Supplemental Figure 6.**
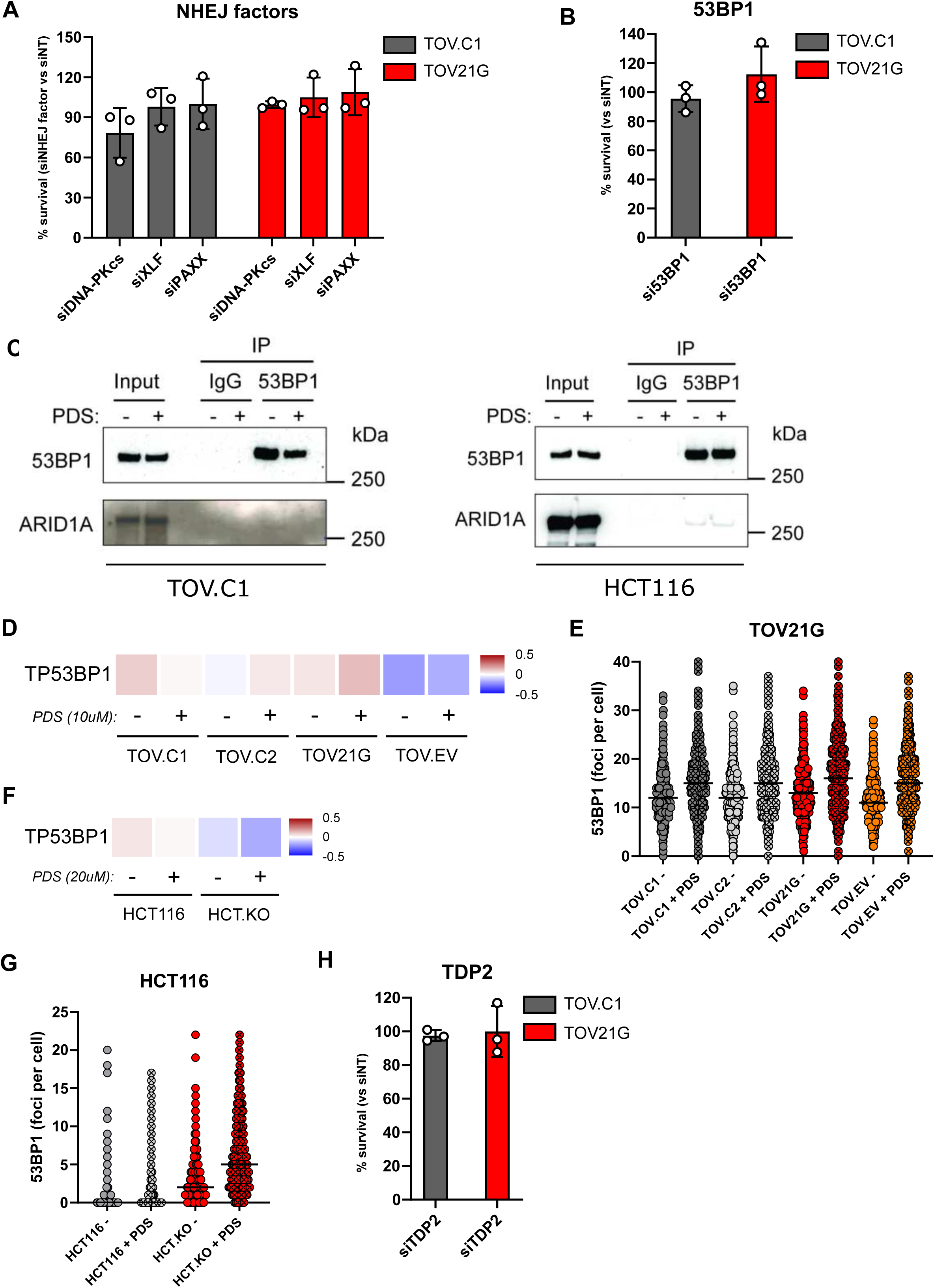
Analysis of 53BP1 and NHEJ factors with ARID1A. (A) Bar graph showing survival of TOV.C1 and TOV21G parental cells upon knockdown of core NHEJ factors, compared to a non-targeting control (siNT), under basal conditions. Points correspond to independent biological replicates, n = 3, mean ± SEM. (B) Bar graph showing survival of TOV.C1 and TOV21G parental cells upon knockdown of 53BP1, compared to siNT, under basal conditions. Points correspond to independent biological replicates, n = 3, mean ± SEM. (C) Co-immunoprecipitation of 53BP1 and ARID1A. 53BP1 was immunoprecipitated from cell extracts prepared from untreated or PDS-treated (10 µM, 4 hours) TOV.C1 cells (left panel) or from untreated or PDS-treated (20 µM, 4 hours) HCT116 cells (right panel). IgG was used as a negative control. (D) Heatmap showing 53BP1 (TP53BP1) enrichment in chromatin in untreated or PDS-treated (10 µM, 4 hours) TOV21G isogenic cell lines. (E) Quantification of 53BP1 foci per cell by immunofluorescence in TOV21G derived cell lines, either untreated or treated with PDS (10 µM, 6 hours). Points represent number of foci in a single nucleus, line represents the median. (F) Heatmap showing 53BP1 (TP53BP1) enrichment in chromatin in untreated and PDS-treated (20 µM, 4 hours) HCT116 parental and ARID1A knockout cells. (G) Quantification of 53BP1 foci per cell by immunofluorescence in HCT116 parental and ARID1A knockout cell lines, either untreated or treated with PDS (20 µM, 6 hours). Points represent number of foci in a single nucleus, line represents the median. (H) Bar graph showing survival of TOV.C1 and TOV21G parental cells upon knockdown of TDP2, compared to siNT, under untreated conditions. Points correspond to independent biological replicates, n = 3, mean ± SEM. *All siRNAs were used in parallel in each of the three biological replicates, hence the non-targeting siRNA controls are the same for each set of siRNAs represented in A, B and H*.

